# Soybean RIN4 represents a mechanistic link between plant immune and symbiotic signaling

**DOI:** 10.1101/2023.09.12.557450

**Authors:** Katalin Tóth, Daewon Kim, Sung-Hwan Cho, Cuong T. Nguyen, Tran H. N. Nguyen, Christopher Hartanto, Jean-Michel Michno, Adrian O. Stec, Robert M. Stupar, Gary Stacey

## Abstract

The legume-rhizobium symbiosis represents a unique and beneficial interaction between legumes and nitrogen-fixing soil bacteria, called rhizobia. The initiation and development of this symbiosis is complex and begins with recognition of key molecular signals, produced by the plant and its symbiont, which determine symbiotic compatibility. Current data suggest that the invading symbiont initially triggers plant immune responses that are subsequently suppressed. Hence, there is growing evidence that features of plant immunity may be relevant to symbiotic establishment. RIN4 is a key immune regulator in plants, regulating basal immunity and it is also targeted by pathogen effector proteins that either confer susceptibility or resistance, depending on the presence of the appropriate resistance protein. Surprisingly, we found that RIN4 was rapidly phosphorylated upon rhizobial inoculation of soybean root hairs. RNAi silencing and mutant studies indicate that RIN4 expression is essential for effective nodulation of soybean. RIN4 phosphorylation occurs within a fifteen amino acid motif, which is highly conserved within the Fabales (legumes) and Rosales orders, which comprise species capable of nitrogen-fixing endosymbiosis with rhizobia. RIN4 proteins mutated in this conserved phosphorylation site failed to support efficient soybean nodulation. Phosphorylation of this site is mediated by the symbiotic receptor-like kinase, SymRK, a well-studied member of the symbiotic signaling pathway. The data implicate RIN4 phosphorylation as a key mediator of rhizobial compatibility, interconnecting symbiotic and immune signaling pathways.

**Significance:** The nitrogen fixing legume-rhizobium symbiosis is a cornerstone of sustainable agriculture, with ongoing efforts to transfer this unique ability to non-leguminous crop plants. Plants are surrounded by a myriad of microbes in the soil, and, therefore, require constant surveillance in order to distinguish between a pathogen or symbiont. Plants monitor for specific molecular signals that indicate pathogen or symbiont presence. We show that RIN4, a key immune regulator, plays an essential role in promoting the development of the symbiotic nitrogen-fixing relationship between soybean and its compatible symbiont *Bradyrhizobium japonicum*. Therefore, RIN4 is likely a key player in mediating the appropriate response upon infection by friend or foe.

## Introduction

At the very beginning of the symbiotic signaling pathway, there are three receptor-like kinases (RLKs) that are indispensable for the initiation of two developmental processes: bacterial infection and nodule organogenesis. Bacterial infection occurs primarily via an infection thread (IT) developed within the infected root hair. IT delivers rhizobia into the underlying newly divided cortical cells forming the nodule primordium, which occurs in parallel with bacterial infection (1; 2/Quilbe et al., 2022). Underneath the infection site, the nodule primordium develops into a new organ, the nodule, where the rhizobia are accommodated and convert atmospheric nitrogen into ammonia. As a result, a nitrogen-fixing symbiosis will develop (1, 2). In legumes with determinate nodules, such as soybean (*Glycine max*) and *Lotus japonicus*, two Lysin (LysM)-domain containing RLKs, Nodulation Factor Receptor 1 and 5 (NFR1 and NFR5) perceive the rhizobial lipo-chitooligosaccharide nodulation factor (NF) (3, 4, 5, 6). Rhizobia produce NF in response to flavonoids secreted by the host legume. Lotus or soybean mutants lacking NFR1 and/or NFR5 do not respond to rhizobial inoculation and do not form nodules (4). A third RLK, containing extracellular leucine-rich repeats, is located downstream of the NF receptors, called Symbiosis Receptor Kinase (SymRK) (7, 8). Root hairs of Lotus *symrk* mutant plants (i.e., cac41.5 insertion mutant) do not curl and bacterial infection cannot occur despite displaying root hair deformation (7). *symrk-10*, a Lotus mutant carrying a point mutation in the activation loop of the kinase domain, abolishing kinase activity, displays a similar phenotype as the insertion mutant (9), underpinning the importance of phosphorylation in the symbiotic signaling cascade.

Studies using Lotus showed that SymRK interacts strongly with NFR5 and weakly with NFR1 (10). It was shown that autophosphorylation of NFR1 is essential for downstream signaling. NFR5 lacks kinase activity and is trans-phosphorylated by NFR1 (11), and by a third LysM-containing RLK, NFRe (epidermal), enhancing the robustness of NF-signaling (12). Transducers of RLK-induced signaling are receptor-like cytoplasmic kinases (RLCKs), a NFR5-interacting cytoplasmic kinase 4 (NiCK4) was shown to be an important link between NF perception by NFR5 and nodule organogenesis (13). Briefly, downstream components of the pathway are a calcium- and calmodulin-dependent kinase, CCaMK which phosphorylates CYCLOPS, a DNA-binding transcriptional activator (14). The CCaMK/CYCLOPS complex controls bacterial infection as well as nodule organogenesis (14), forming a regulatory unit with other transcriptional regulators and activates NODULE INCEPTION (NIN), a nodulation specific transcription factor. NIN is involved in root hair and epidermal as well as cortical cell responses, the latter leading to nodule development (15).

Given the importance of phosphorylation in the symbiotic signaling cascade, it is not surprising that phosphoproteomic studies have been reported for a variety of legumes (16, 17, 18). The primary entry point for rhizobium in the case of *L. japonicus*, *Medicago truncatula*, and *G. max* is the root hair (19). However, only a small fraction of the root hairs on a given root are infected and even fewer infections lead to nodule formation (20). Hence, phosphoproteomic studies using entire roots, such as conducted with *L. japonicus* and *M. truncatula,* likely suffer from signal dilution due to the highly localized nature of rhizobial infection. Therefore, we previously performed phosphoproteomic studies of isolated soybean root hair cells (separated from the root), in order to reduce signal dilution due to non-responding root tissues (17). Indeed, this study identified a variety of proteins that were rapidly (one-hour post-inoculation) phosphorylated upon treatment with the compatible symbiont, *Bradyrhizobium japonicum.* To our surprise, among these proteins was the plant immune regulator RPM1-INteracting protein 4 (RIN4; 17).

RIN4 was discovered in *Arabidopsis thaliana* as an interactor of RPM1, a disease resistance protein conferring resistance against the bacterial leaf pathogen *Pseudomonas syringae* (21). RIN4 is conserved among land plants and is involved in the regulation of Pattern Triggered Immunity (PTI). Given RIN4’s regulatory function in PTI, it is not surprising that the protein is targeted by several effector proteins released by pathogens to interfere and modulate plant immune responses (22). RIN4 undergoes post-translational modifications (PTM) or proteolytic cleavage as a consequence of being targeted by *P. syringae* effector proteins (23, 24). RIN4 modifications trigger a second layer of immune responses triggered by resistance (R) proteins, intracellular immune receptors (nucleotide-binding leucine-rich repeat receptor or NLRs) which monitor perturbations within the host cell leading to NLR-triggered immunity (NTI) (24). RIN4 is an intrinsically disordered protein (IDP) (25, 26). IDPs lack stable secondary and tertiary protein structure and can transition from disorder to order upon interactions with other protein(s) or upon PTMs like phosphorylation (27). Lee and colleagues (26) demonstrated that phosphorylated RIN4 is more flexible than native RIN4 contributing to its conformational flexibility and function.

*P. syringae* effector proteins AvrB and AvrRpm1 induce RIN4 phosphorylation, suppressing PTI responses (28, 29). Phosphorylation of AtRIN4 at serine 141 is triggered upon bacterial flagellin treatment (i.e., flagellin epitope, flg22). Phosphorylation of this site is the target of AvrB and consequently induces phosphorylation of the evolutionary conserved threonine 166. Increased T166 phosphorylation suppresses S141 induced PTI responses (29). It was shown that AvrRpm1 ADP-ribosylates Arabidopsis as well as soybean RIN4, and this ADP-ribosylation is a prerequisite for subsequent phosphorylation of the T166 phosphorylation site (30). AvrB-induced T166 phosphorylation is mediated by an RLCK, RIPK. However, in an Arabidopsis *ripk* mutant background, RIN4 phosphorylation in response to AvrB is decreased and not abolished (31). Furthermore, Xu and colleagues (32) showed that several other RLCKs were able to phosphorylate RIN4. Hence, the relative phosphorylation of S141 and T166 determines the plant response to pathogen effector proteins and subsequent disease progression.

Although RIN4 plays an essential role in pathogen virulence and host immunity, the details of RIN4 molecular function are not well understood (24). It should be noted that virtually all the studies on RIN4 have used leaves, the natural infection route for *P. syringae*.

The *A. thaliana* genome encodes a single *RIN4* gene. In contrast, the soybean genome encodes four *RIN4* (*GmRIN4a-d*) genes (33). In soybean, RPG1-B (resistance to *Pseudomonas syringae pv glycinea*) R protein conveys resistance to *P. syringae* expressing AvrB. Both GmRIN4a and GmRIN4b were shown to associate with AvrB, but only GmRIN4b interacts with RPG1-B (33). These data and the above mentioned AvrRpm1-mediated ADP-ribosylation of both At and GmRIN4 suggest that, at least in leaves, the GmRIN4 proteins play a role in plant immunity similar to that defined by detailed studies in *Arabidopsis*.

In the work described here, we demonstrate that soybean RIN4 (GmRIN4, hereafter RIN4) protein(s) are essential for efficient nodulation of soybean. This function of RIN4 is mediated by specific phosphorylation of serine 143, which is located within a 15 amino acid (aa) motif. These 15 aa are absent in *Arabidopsis*, and seem to be highly conserved within the Fabales and Rosales plant orders, therefore suggesting a symbiosis-related function. Phosphorylation of S143 is mediated by GmSymRKβ. We found that *RIN4a* and *RIN4b* are highly expressed in root hair cells and that their expression level does not change upon rhizobial treatment. A soybean mutant line, in which *RIN4b* was mutated using CRISPR-Cas9 mediated gene editing, led to a significant reduction in nodulation concomitant with the reduction in the expression of downstream components of the symbiosis signaling pathway.

## Results and Discussion

### Soybean RIN4 proteins harbor a highly conserved and novel-RIN4-motif

A key, unifying feature of the nitrogen-fixing symbiosis is the formation of nodules where the bacteria are accommodated inside living plant cells (34). The symbiosis is restricted to one phylogenetic clade containing four orders: Fabales (legumes), Fagales, Cucurbitales and Rosales (FaFaCuRo). Within this clade, there are 10 families out of the 28 that contain species which form nitrogen-fixing root nodules (34). Legumes and the non-legume *Parasponia andersonii* (Rosales) form symbiosis with rhizobia. Actinobacteria, *Frankia*, interact with species from Rosales, Fagales and Cucurbitales (35), which also leads to intracellular symbiotic nitrogen fixation.

We built a phylogenetic tree (SI Appendix, Fig. S1 and Fig. 1A) of 149 RIN4 sequences derived from 66 species of the FaFaCuRo clade and species outside of the clade (Materials and Methods and SI Appendix, Table S1). The tree contains sequences from both nodulating and non-nodulating species of Fabales, Rosales, Fagales and Cucurbitales (35), as well as from species outside of the clade. This analysis identified a specific clade of RIN4 proteins (SI Appendix, Fig. S1). Two subclades were apparent: one of them comprised of sequences from Fabales (SI Appendix, Fig. S1, blue highlighted), the other contains sequences from Rosales (SI Appendix, Fig. S1, green highlighted). Within the sequences forming these two apparent subclades, we discovered a 15 amino acid motif, defined by a “GRDSP” core sequence (Fig. 1B, red box), suggesting a nodulation related function. We named it a novel-RIN4-motif (NRM) (Figure 1B, C). In Figure 1, the alignment shows RIN4 protein sequences from model legume species such as *G. max*, *P. vulgaris*, *L. japonicus* and *M. truncatula* aligned with *Arabidopsis* RIN4, and nodulating non-legume *P. andersonii* and its non-nodulating relative *Trema orientale* from Rosales. The sequence is absent in *Arabidopsis* RIN4 (Fig. 1B, red box) as well as in other non-FaFaCuRo species we used when building the phylogenetic tree. This motif is highly conserved among RIN4 proteins from nodulating and non-nodulating species of Fabales as well as Rosales. Interestingly, the motif is not conserved in RIN4 sequences derived from Fagales and Cucurbitales (SI Appendix, Fig S2). However, the motif is retained in non nodulating Fabales (such as *C. canadensis*, *N. schottii*; SI Appendix, Fig. S2) and Rosales species as well. NRM harbors the soybean RIN4 phosphorylation site, serine 143 (Fig. 1B, C) identified in our previous study of soybean root hairs (17). S143 is localized within the “GRDSP” core motif (Fig. 1B, red arrow), and is highly conserved across Fabales and Rosales.

**Fig. 1.**
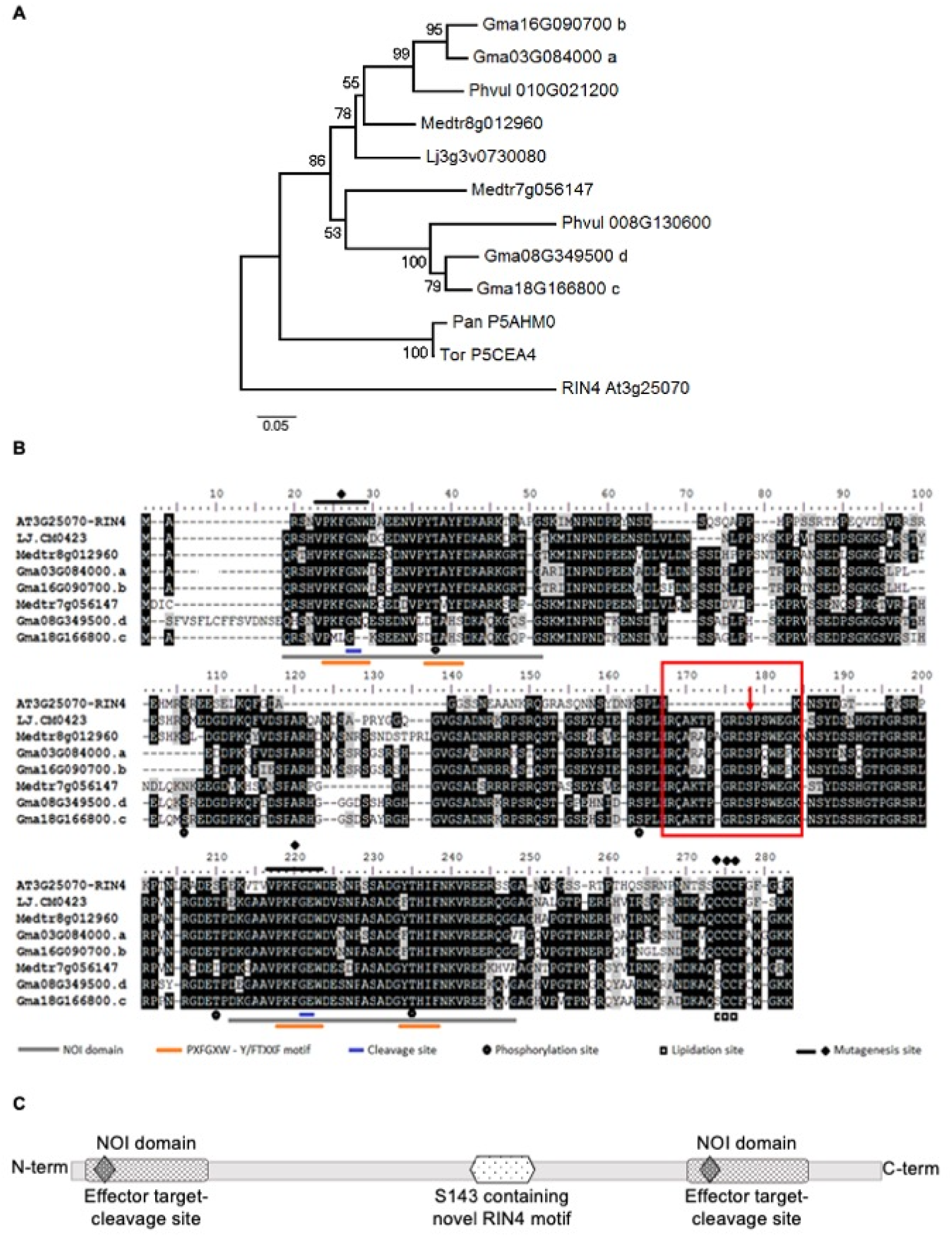
Phylogeny of RIN4 protein homologs from model legume species and a novel-RIN4- motif within soybean RIN4. (A) Phylogeny of soybean RIN4 closest homologs including GmRIN4a, GmRIN4b, GmRIN4c, GmRIN4d, *L. japonicus* RIN4 (Lj3g3v0730080), two *M. truncatula* RIN4 (Mtr8g012960, Mtr7g056147), two *P. vulgaris* RIN4 (Pvul010G021200, Pvul008G130600), and nodulating non-legume *P. andersonii* (Pan P5HM0) and its non-nodulating relative *Trema orientale* (Tor P5CEA4). Tree was rooted using *A. thaliana* RIN4 (At3g25070). (B) Novel-RIN4-motif within RIN4 protein sequences (red box). One of the identified phosphorylation sites, S143 is located within this motif (red arrow) within a “GRDSP” core sequence that is highly conserved among legume species and species of Rosales. Here, we show the sequence alignment of *Arabidopsis* RIN4 with soybean (Gma), *L. japonicus* (Lj), *M. truncatula* (Mtr), *P. vulgaris* (Pvul) and nodulating non-legume *P. andersonii* (Pan) and its non-nodulating relative *T. orientale* (Tor) RIN4 proteins. Grey underline indicates Nitrate-induced domain (NOI). Red underline indicates motif for proteolytic cleavage targeted by pathogenic effector proteins, while blue line shows cleavage site. The characteristic feature of the NOI-domain is that it harbors the PXFGXW motif which is the target site for effector protein. Tryptophan (W) is a crucial residue within the motif. RIN4c and RIN4d are missing W (blue arrow), and SMART analysis also predicted only one Pfam: AvrPt2 cleavage site. (C) RIN4 protein schematic based on soybean RIN4a/b. NOI domains at the proteins N- and C-terminus, diamond within NOI domain represents cleavage site targeted by effector protein(s), hexagon represents the Novel-RIN4-motif containing S143 phosphorylation site.

### *RIN4a* and *RIN4b* are highly expressed in soybean root hairs

There are four *RIN4* genes described in soybean (*RIN4a, RIN4b, RIN4c, RIN4d*; 33; Fig. 1). A characteristic feature of the AtRIN4 protein is two plant-specific nitrate-induced domains (NOI), an N-terminal and a C-terminal domain (Fig. 1B, grey underline; Fig 1C). AvrRpt2 bacterial effector targeted cleavage sites are located within these motifs (36, Fig. 1B, red underline; Fig 1C). Our *in-silico* analysis found that all four soybean RIN4 proteins contain 2 NOI domains, whereas the AvrRpt2 cleavage site is absent in RIN4c and RIN4d N-terminal NOI domains (Fig. 1B, blue arrow). This observation is based on sequence alignment (Fig. 1B) and prediction using the Simple Modular Architecture Research Tool (SMART).

Root hairs are the primary entry point for rhizobial infection in most legumes (2). We wanted to narrow the number of study subjects and, therefore, looked at the gene expression level of *RIN4* genes in soybean root hairs and roots using quantitative reverse transcriptase polymerase chain reaction (qRT-PCR) analysis. *RIN4a* and *RIN4b* showed much higher expression in root hairs than *RIN4c*, *RIN4d* and a *RIN4-like* gene (SI Appendix, Fig. S3A). *RIN4* transcripts levels were not induced upon rhizobial inoculation (SI Appendix, Fig. S3A), suggesting that the protein is regulated post-translationally, as was previously observed in other studies. In stripped roots (roots with root hairs removed), all *RIN4* genes displayed lower expression in comparison to root hairs, and none were up-regulated upon rhizobial treatment (SI Appendix, Fig. S3A). These results are consistent with *Arabidopsis* RNA-seq data revealing that *AtRIN4* is one of the most abundant transcripts in root hairs (37). Given that plant roots are constantly surrounded with microbes in the rhizosphere, it is not surprising that such a key immune regulator as RIN4 is being expressed and maintained at a high level. Our gene expression data were confirmed by Western-blot analysis using total protein extracted from root hairs and stripped roots (SI Appendix, Fig. S3B). RIN4 was detected using custom-specific anti RIN4 antibody generated against a mixture of RIN4a and RIN4b recombinantly expressed proteins. Anti-RIN4 antibody was tested using His-epitope tagged RIN4a, RIN4b, RIN4c and RIN4d recombinantly expressed and purified proteins. The RIN4 antibody recognizes only RIN4a and RIN4b, and not RIN4c and RIN4d (SI Appendix, Fig. S3C).

RIN4a and RIN4b share 93% identity on an amino acid level, whilst RIN4a/b share 64% identity with RIN4c and 62% identity with RIN4d on an amino acid level. RIN4c and RIN4d displayed lower expression in root hairs and roots compared to RIN4a and RIN4b (SI Appendix, Fig. S3). Furthermore, it was previously shown that only *RIN4b* complements *AtRIN4* in an *Arabidopsis rin4* mutant background (33). Therefore, we focused our attention on RIN4a and RIN4b in this study.

### *RIN4a* and *RIN4b* are required for efficient nodulation

Relatively few soybean mutants are available and, therefore, we searched for *rin4* mutants in the model legume *L. japonicus,* which forms determinate nodules, as does soybean. In *L. japonicus,* we could identify only one *RIN4* gene (Fig. 1). From the *L. japonicus* LORE1 mutant population database (https://lotus.au.dk/; 38, 39), we ordered two lines with exogenic LORE1 insertions in the *RIN4* gene locus (Plant ID 30000711, 30019656). Genotyping of the two exogenic lines did not reveal homozygous individuals (SI Appendix, Table S2), suggesting that *RIN4* homozygous mutation might be lethal in *L. japonicus*, as is the case in *A. thaliana* (21).

In order to assess the role of *RIN4* genes during the legume-rhizobium symbiosis, we targeted *RIN4a* and *RIN4b* by RNAi-mediated gene silencing. *RIN4a* and *RIN4b*-RNAi constructs were introduced into soybean roots via Agrobacterium-mediated hairy root transformation. Silencing of *RIN4a* and *RIN4b* resulted in significantly reduced nodule numbers (about 50-70% less) on soybean transgenic roots in comparison to transgenic roots carrying the empty vector control (Fig. 2A and B). Based on the qRT-PCR data (Fig. 2C), the transcript levels of both genes were significantly reduced, but not abolished, suggesting that both genes contribute to the formation of the symbiosis. Transcripts of the two genes are 92% identical, explaining the reduction in both transcripts (Fig. 2C). Because of the high-level identity of the two genes, it was very challenging to silence each gene separately.

**Fig. 2.**
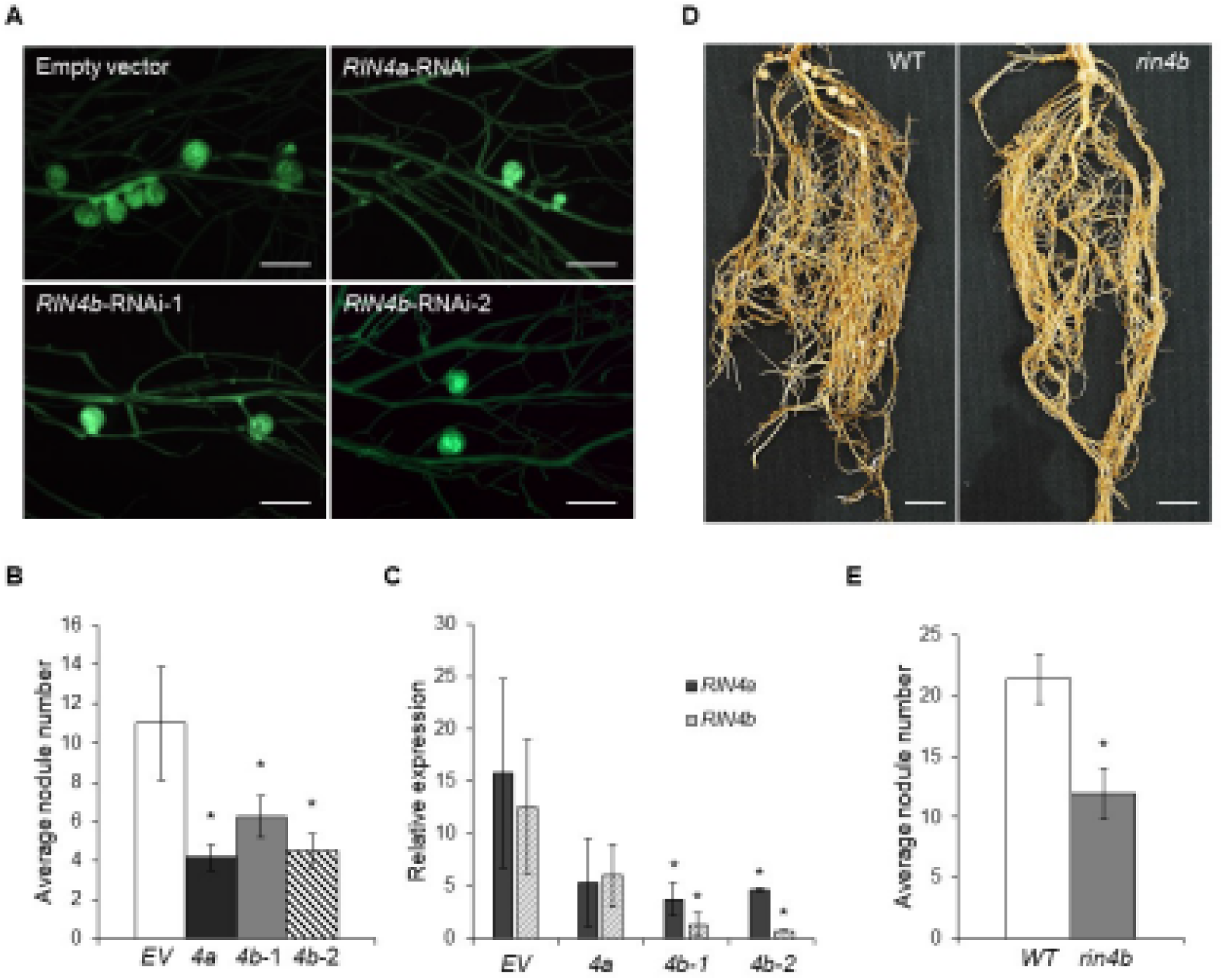
*RIN4a* and *RIN4b* are required for proper symbiosis formation. (A) Micrographs showing representative transgenic roots of gene silencing, visualized by green fluorescence originating from GFP marker carried by all vectors. Scale bars represent 2 mm. (B) Significantly reduced nodule numbers were observed on soybean transgenic roots carrying RNAi constructs targeting *RIN4a* and *RIN4b*. Roots were phenotyped 5 wpi. Representative data of one biological replicate, experiment was done in 3 biological replicates. Student’s t-test * p<0.05. (C) Quantitative reverse transcription polymerase chain reaction (qRT-PCR) analysis confirmed reduced transcript levels of *RIN4a* and *RIN4b* in RNAi transgenic roots, strongly suggesting that the individual constructs targeted both genes. (D) Roots of wild-type *RIN4b* (WT) and mutant *rin4b* (in Bert background) homozygous mutant plants. Scale bars represent 2 cm. (E) Reduced nodule numbers in Bert soybean roots carrying a CRISPR-Cas9 edited 2 bp deletion in *RIN4b* (*rin4b*) in comparison to plants expressing wild-type *RIN4b* (WT) which out segregated as non-transgenic and non-edited. Experiments were done twice. Student’s t-test * p<0.05.

To further confirm that soybean *RIN4a* and *RIN4b* play a role in formation of the nitrogen-fixing symbiosis, we targeted both genes using CRISPR-Cas9 gene editing technology to generate stable knock-out mutants. A CRISPR-Cas9 edited soybean knock-out line was obtained only in *RIN4b* in the Bert cultivar background (40). This line contains a two base pair deletion within the second exon of the gene that results in a premature stop codon leading to reduced RIN4 protein abundance and significantly reduced *RIN4b* mRNA levels (SI Appendix, Fig. S4). This line produced significantly reduced nodule numbers in comparison to plants expressing wild-type *RIN4b* (Fig. 2D and E), supporting the findings of RNAi-mediated knock down and further confirming the role of *RIN4b* in the symbiosis. Hand-made cross-sections revealed that nodules on both wild-type and *rin4b* mutant roots were pink inside. The pink color reflects the presence of leghemoglobin, suggesting that the nodules were functional and fixing nitrogen. *rin4b-CRISPR-Cas9* produced about 50-60% less nodules suggesting that there may be functional redundancy between *RIN4a* and *RIN4b*.

In our previous phosphoproteomic study, we found that RIN4 protein(s) phosphorylation occurred one-hour post-inoculation (hpi) (16). One of the identified phosphorylation sites is S143 (SI Appendix Fig. S5A). Intriguingly, S143 is harbored within the highly conserved “GRDSP” core motif located in the NRM (Fig. 1B, red arrow). The NRM is 100% identical between RIN4a and RIN4b (Fig. 1B). Given that the phosphorylation site is located within the NRM, we decided to further investigate the function of the S143 phosphorylation site. A pS143-specific peptide antibody was generated to detect phosphorylation (SI Appendix, Fig. S5A). We treated three-day old soybean seedlings with mock (H2O) and wild-type *B. japonicum*, to confirm the phosphorylation previously observed. Root hairs (RH) were harvested 1 hpi separately from root tissue. AtRIN4 displays a basal phosphorylation level in mock-treated plants (26, 29); therefore, it is not surprising that our pS143 specific antibody shows RIN4 phosphorylation in mock-treated RH (SI Appendix Fig. S5B). ImageJ quantification of phosphorylation showed a 2-fold upregulation in response to *B. japonicum* when compared to mock-treated RH (SI Appendix Fig. S5C). To address the function of the identified S143 phosphorylation site in relation to the symbiosis, we introduced point mutation(s) by site-directed mutagenesis and generated RIN4a^S143A^ and RIN4b^S143A^ mutated proteins that cannot be phosphorylated at S143. Aspartic acid (D) was introduced to mimic the phosphorylation status and phospho-mimic mutant versions (RIN4a^S143D^ and RIN4b^S143D^) were created. RIN4a and RIN4b were N-terminally tagged with HA-epitope to detect the presence of the introduced mutant and native versions of the protein in soybean transgenic roots. Interestingly, ectopic expression of RIN4a^S143A^ and RIN4b^S143A^ mutant proteins led to significantly reduced nodule numbers on transgenic roots in comparison to roots expressing wild-type RIN4b and empty vector control, whereas expression of RIN4a^S143D^ and RIN4b^S143D^ did not affect nodule numbers (SI Appendix, Fig. S6). These results suggest that phosphorylation at the S143 site in RIN4a and RIN4b is required for efficient symbiosis formation. In order to verify protein expression in transgenic roots, Western-blot analysis was performed on total protein extracts from transgenic roots. All constructs displayed two protein bands (phosphorylated and non-phosphorylated versions) at the expected size (around 30 kDa) showing that the introduced versions of RIN4a and RIN4b were expressed (SI Appendix, Fig. S6), contributing to the observed phenotype.

### Phosphor-negative RIN4b^S143A^ does not complement *rin4b* mutant

Complementation of the *rin4b*-CRISPR-Cas9 mutant was carried out by introducing either RIN4b, RIN4b^S143A^, RIN4b^S143D^ or empty vector (control) via Agrobacterium-mediated hairy root transformation into soybean transgenic roots induced on *rin4b* mutant plants. This experiment confirmed that RIN4bS143 is critical for RIN4b symbiotic function, as the phosphor-negative mutant version (RIN4b^S143A^) of the protein, similar to the empty vector, was unable to rescue the nodulation phenotype observed on the mutant plants (Fig. 3A and B). In contrast, transgenic roots expressing RIN4b or the phosphomimic version RIN4b^S143D^ restored nodulation in comparison to transgenic roots carrying empty vector (Fig. 3). Therefore,we refer to S143 as nodulation-related S143 in the following parts of the manuscript. Expression of the HA-tagged RIN4b wild-type and mutant proteins in the transgenic roots was confirmed via Western-blot analysis (Fig. 3C).

**Fig. 3.**
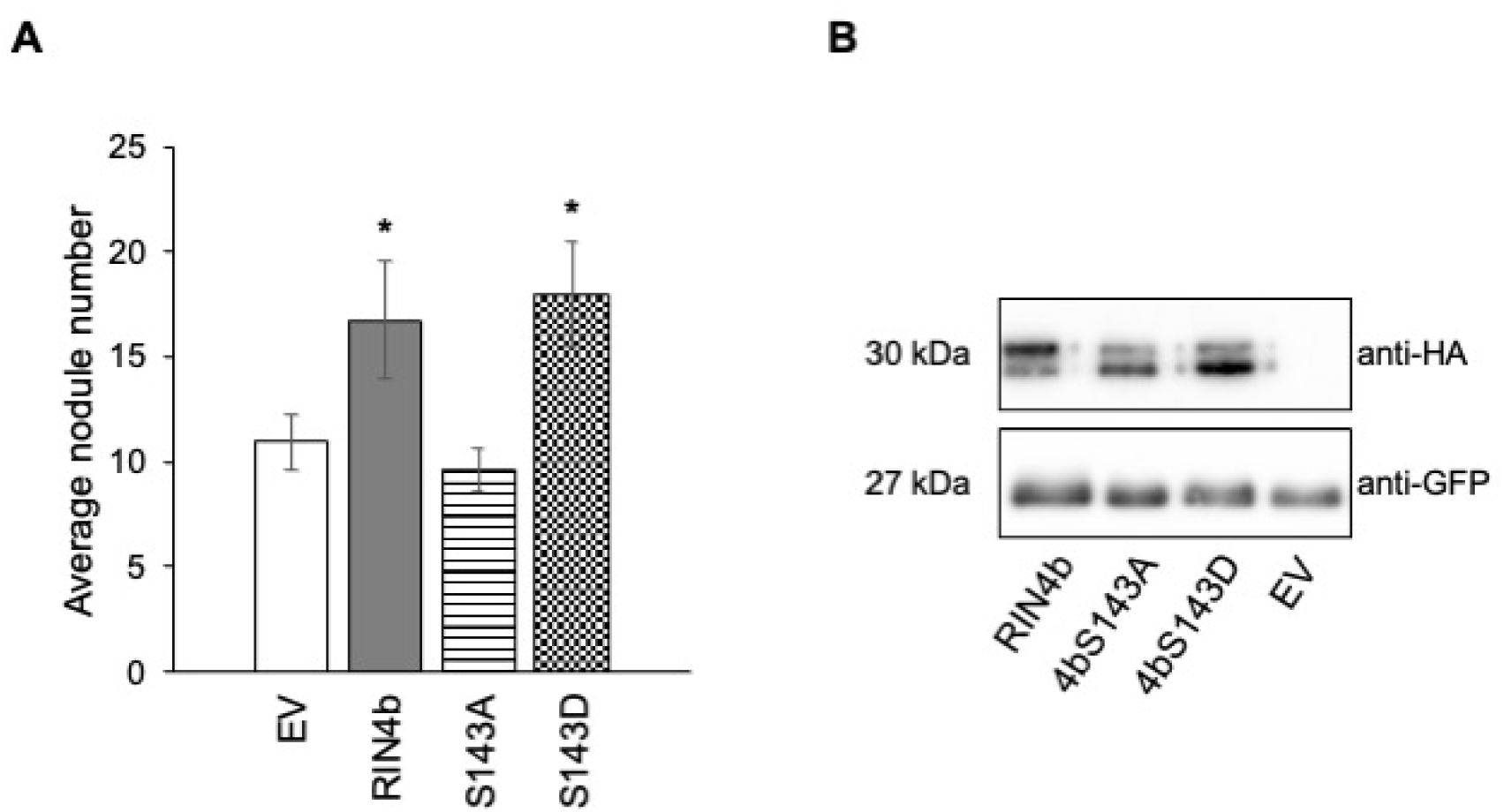
Phosphor-negative RIN4b^S143A^ does not complement *rin4b* mutant phenotype. (A) *rin4b*-CRISPR-Cas9 mutant was transformed with RIN4b, RIN4b^S143A^, RIN4b^S143D^ and empty vector. Only RIN4b and RIN4b^S143D^ could rescue the phenotype caused by *rin4b* mutation (micrographs on the right). RIN4b^S143A^ and an empty vector could not restore nodule numbers on the transgenic roots (micrographs on the left). Micrographs showing representative transgenic roots expressing the respective constructs, visualized by green fluorescence originating from GFP marker. Scale bar represents 1 mm. (B) Graphical representation of the results from 2 biological replicates. Error bars represent standard error. Student t-test, * p<0.05. (C) Western-blot analysis showed that the transgenic roots expressed the HA-tagged RIN4b proteins, as well as the GFP marker carried by the vector.

### RIN4a and RIN4b closely associate with symbiotic receptor-like kinases NFR1α and SymRKß *in planta*

There are two active kinases required for early signal transduction during legume symbiosis development, NFR1 and SYMRK (11, 9). In soybean, both NFR1 and SymRK are present in two copies (NFR1α/β and SymRKα/β; 6, 8). Ectopic over-expression of NFR1α led to increased nodule numbers on transgenic roots, whereas this phenotype was not observed when NFR1β was overexpressed (6). RNAi-mediated gene silencing was performed on both soybean *SymRK* genes. Silencing of *SymRKβ* showed a stronger phenotype suggesting that this protein has the major function in nodulation (8). Since NFR1α and SymRKβ seem to be the major players in soybean, we decided to investigate the interaction of RIN4a and RIN4b only with NFR1α and SymRKβ and not their homologs (NFR1β and SymRKα). Previously it was shown that SymRK in *L. japonicus* undergoes proteolytic cleavage, when the Malectin-like domain (MLD) within the protein’s extracellular region is cleaved off creating SymRKΔMLD (10). ΔMLD was easier to detect when expressing both full-length (FL) SymRKß and SymRKßΔMLD in tobacco leaves. Therefore, for future experiments we used the SymRKßΔMLD construct, instead of SymRKß-FL. In the *in planta* Bimolecular Fluorescence Complementation assay (BiFC), RIN4a and RIN4b showed interaction with each other as previously shown (33) and were used as a positive control for the assay (SI Appendix, Fig. S8A). Co-expression of RIN4a/b with SymRKßΔMLD as well as NFR1α resulted in YFP fluorescence detected by Confocal Laser Scanning Microscopy (SI Appendix, Fig. S8) suggesting that RIN4a and b proteins are closely associated with both RLKs. No interaction could be observed between RIN4a/b and the P2K1 receptor, which was used as negative control in the BiFC assay (SI Appendix, Fig. S8 D and G). The reconstituted YFP signal localizes to the plasma membrane (PM) in accordance with earlier observations where the investigated proteins were localized to the PM (10, 11, 33). PM was visualized by application of a membrane staining dye, FM4-64 (SI Appendix, Fig. S8). The interaction between RIN4a/b and NFR1α and SymRKßΔMLD was confirmed in a Split-Luciferase Complementation assay (SI appendix, Fig. S9), when the proteins were fused to the N-terminal and C-terminal domains of the Luciferase enzyme, which upon reconstitution results in bioluminescence.

### SymRKβ phosphorylates the nodulation-related S143 phospho-site *in vitro* and *in planta*

The kinase activity of soybean NFR1α was demonstrated by Choudhury and Pandey (41). Here, we show that soybean SymRKβ possesses an active kinase domain (SymRKß KD, SI Appendix, Fig. S10). The isolated soybean SymRKβ-KD showed strong autophosphorylation activity, as well as the ability to trans-phosphorylate MBP, a universal substrate (SI Appendix, Fig. S10). Furthermore, to confirm that phosphorylation was triggered by SymRKβ-KD, we introduced a point mutation in the same position as previously described by Yoshida and Parniske (41) for the *L. japonicus* SymRK kinase domain. Specifically, D734N (which corresponds to D738 in LjSymRK) was introduced, in the activation loop, and led to inactivation of the SymRKβ kinase activity (SI Appendix, Fig. S10).

Both NFR1α and SymRKβ kinase domains phosphorylated RIN4a and RIN4b *in vitro* when radioactive [γ-32P] ATP was used to visualize the phosphorylation (SI Appendix, Fig. S11 A).

In order to ascertain which of these two kinases phosphorylates the nodulation-related RIN4S143, we performed mass spectrometry-based Absolute Quantification (AQUA), a method that uses stable-isotope labeled peptides as internal standards to quantify proteins or post-translational modifications (43). The abundance of the heavy-labeled peptides and their corresponding endogenous peptides (peptides derived from native RIN4a and b) can be quantified using selected reaction monitoring mass spectrometry (MS-SRM). In our phosphoproteomic study of soybean root hairs, we also observed phosphorylation of RIN4T173 (17) which corresponds to the earlier published AtRIN4S141 triggered by bacterial flagellum epitope, flg22 (29). Heavy-labeled AQUA peptides were generated against native peptides carrying S143 and T173 (to serve as a control), as well as phosphorylated versions of the peptides (SI Appendix, Table S3).

*In vitro* kinase assay was performed in the absence and presence of ATP (the donor of phosphate group), and MS-SRM was carried out to quantify phosphorylation levels of RIN4a^S143^, RIN4b^S143^, RIN4a^T173^ and RIN4b^T173^. The nodulation-related S143 site was phosphorylated only by SymRKβ in the presence of ATP in RIN4a, as well as in RIN4b (SI Appendix, Fig. S11 B), whereas phosphorylation of RIN4a^T173^ and RIN4b^T173^ was not detected (SI Appendix, Fig. S11 B). No phosphorylation of RIN4a^S143^, RIN4b^S143^, RIN4a^T173^ and RIN4b^T173^ was observed when the proteins were co-incubated with NFR1α-KD (SI Appendix, Fig. S11 B). Calibration curves were established and correlation coefficients were determined for both S143 and T173 peptides (SI Appendix Fig. S12). Peptides in both proteins were detected at a similar level when equal amounts were injected into the mass spectrometer (SI Appendix Fig. S12).

To further assess RIN4a and RIN4b phosphorylation by SymRKβ in an *in planta* environment, the proteins were co-expressed in Arabidopsis leaf protoplasts. Given that AtRIN4 lacks not only the S143 phosphorylation site but also the motif that carries this phosphorylation site, it provided clear evidence about the phosphorylation status of the protein induced by SymRKβ. RIN4a and RIN4b phosphorylation by wild-type SymRKβΔMLD is demonstrated on Figure 4. SymRKβΔMLD kinase inactive version does not phosphorylate either RIN4a or RIN4b. Furthermore, it is also shown that phosphor-minus versions RIN4a^S143A^ and RIN4b^S143A^ do not display a phosphorylated band (Fig. 4), supporting the specificity of the α-pS143 phosphor-specific peptide antibody.

**Fig. 4.**
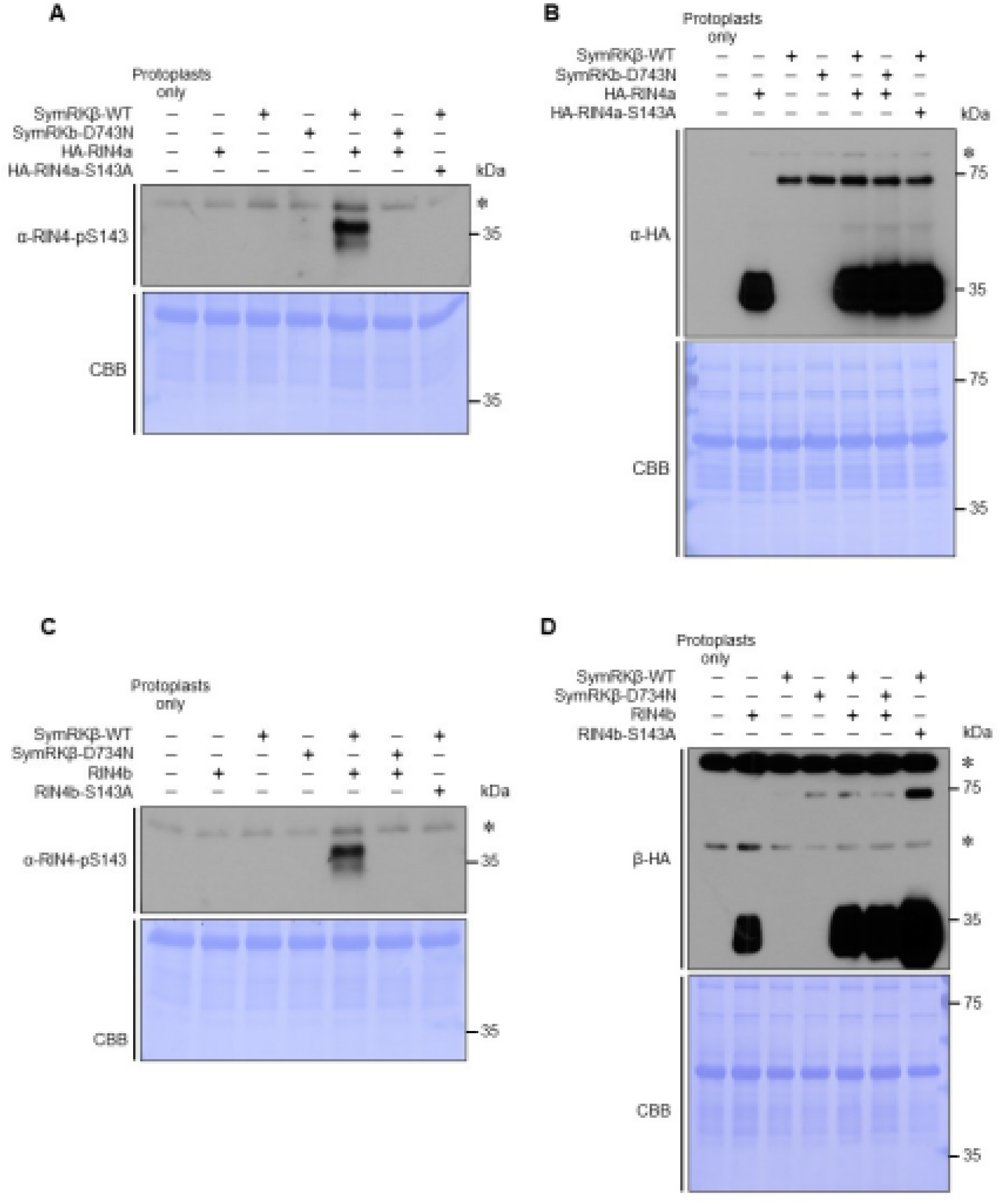
Nodulation-related S143 is phosphorylated by SymRKβ *in planta*. (A) Phosphorylation of RIN4a by SymrkβΔMLD is demonstrated using α-pRIN4-S143, while no phosphorylation can be observed when RIN4a was co-expressed with the kinase inactive version of SymRKβ *in planta*, in *Arabidopsis* protoplasts. (B) shows the expression of all components co-expressed in protoplasts: HA-tagged RIN4a, HA-RIN4aS143A, HA-tagged wild-type SymrkβΔMLD and kinase-dead SymrkβΔMLD. (C) Phosphorylation of RIN4b by SymrkβΔMLD is demonstrated using α-pRIN4-S143, while no phosphorylation can be observed when RIN4b was co-expressed with the kinase inactive version of SymRKβ (D) shows the expression of all components co-expressed in protoplasts: HA-RIN4b, HA RIN4bS143A, wild-type HA-SymrkβΔMLD and HA-SymrkβΔMLD-D734N. *: non-specific band shows equal loading of the protein extracts as well as CBB staining of the membranes displays equal protein extract loading.

### RIN4b acts at the intersection of symbiotic and immune signaling

In order to decipher the RIN4b contribution to symbiotic signaling, we performed gene expression analysis of well-known symbiotic signaling genes, such as the transcription factors (TF) NIN, NF-YA and ERN1 in the *rin4b* (CRISPR-Cas9) mutant background. *NIN* is a nodulation-specific TF, which was the first gene identified in the symbiotic pathway more than 20 years ago (44). Both root epidermal and cortical signaling leads to activation of *NIN*. NIN contributes to bacterial infection in root hairs, to epidermal responses and cortical cell division leading to nodule organogenesis (15). NIN activates Nuclear Factor-Y (NF-Y) transcriptional subunits, a heterotrimeric DNA-binding protein complex composed of NF-YA, NF-YB and NF-YC (15). Soyano and colleagues (15) identified *NF-YA* and *NF-YB* in *L. japonicus* as a target of NIN regulation. *LjNF-YA* overexpression caused changes in the root architecture, while overexpression of *LjNF-YB* did not show root alterations, therefore *LjNF-YA* was designated as the primary player in cortical cell division (15). *ERN1* in *L. japonicus* and *M. truncatula* encodes an AP2/ERF transcription factor. ERN1 is the central regulator of the infection process and is directly regulated by the CCaMK-CYCLOPS complex (45).

Hayashi and colleagues (46) identified four *NIN*-like genes in soybean (*GmNIN1a*, *GmNIN1b*, *GmNIN2a* and *GmNIN2b*). Since *NIN1b* was detected at a low level and did not display significant induction in response to rhizobia (46), only *GmNIN1a*, *GmNIN2a* and *GmNIN2b* were included in our analysis. While it was shown that both *GmERN1a* and *GmERN1b* responded to rhizobial inoculation, in our hands only *ERN1a* could be amplified. Sequence alignment identified at least three NF-YA transcription factor homologs (NF-YA 1, 2, 3) in soybean. Based on preliminary experiments, only *NF-YA1* and *NF-YA3* were found to be expressed and, therefore, *NF-YA2* was excluded from the analysis. For these experiments, *rin4b*-CRISPR-Cas 9 mutant in the Bert background (M4 and M5 bulk) and Bert wild-type seedlings were used. Seedlings four days post-germination were treated with mock (H2O) and *B. japonicum*. Given the limited number of *rin4b* mutant seeds, root hair separation from roots was not possible, therefore entire roots were harvested one hour and 12 hours post-inoculation. *NIN1a* expression (the closest homolog to *LjNIN* and *MtNIN*; 46), in agreement with previously published reports, was induced in wild-type plants 12 hours post-inoculation with

*B. japonicum* (Fig. 5), whereas its expression in *rin4b* mutant was not induced. *NIN2a* and *NIN2b* showed induction in *rin4b* upon rhizobial treatment, but at a significantly lower level in comparison to wild-type roots (Fig. 5). Since NIN is responsible for induction of NF-YAs, their reduced expression in *rin4b* plants was not unexpected (Fig. 5). *ERN1* expression is also affected in *rin4b* mutant plants, this data suggest that RIN4b is located upstream of the CCaMK-CYCLOPS complex, as *ERN1* expression is CYCLOPS dependent (Kawaharada et al, 2017/47) Based on the results of the expression data presented above, it seems that both *NIN*-dependent and *NIN*-independent branches of the symbiotic signalling pathways are impacted in the *rinb4* mutant background. This indicates that the role of RIN4 protein in the symbiotic pathway can be placed upstream of CYCLOPS, as the CYCLOPS transcription activating complex is responsible (directly or indirectly) for the activation of all TFs tested in our expression analysis (Singh et al, 2014/48, 47) .

**Fig. 5.**
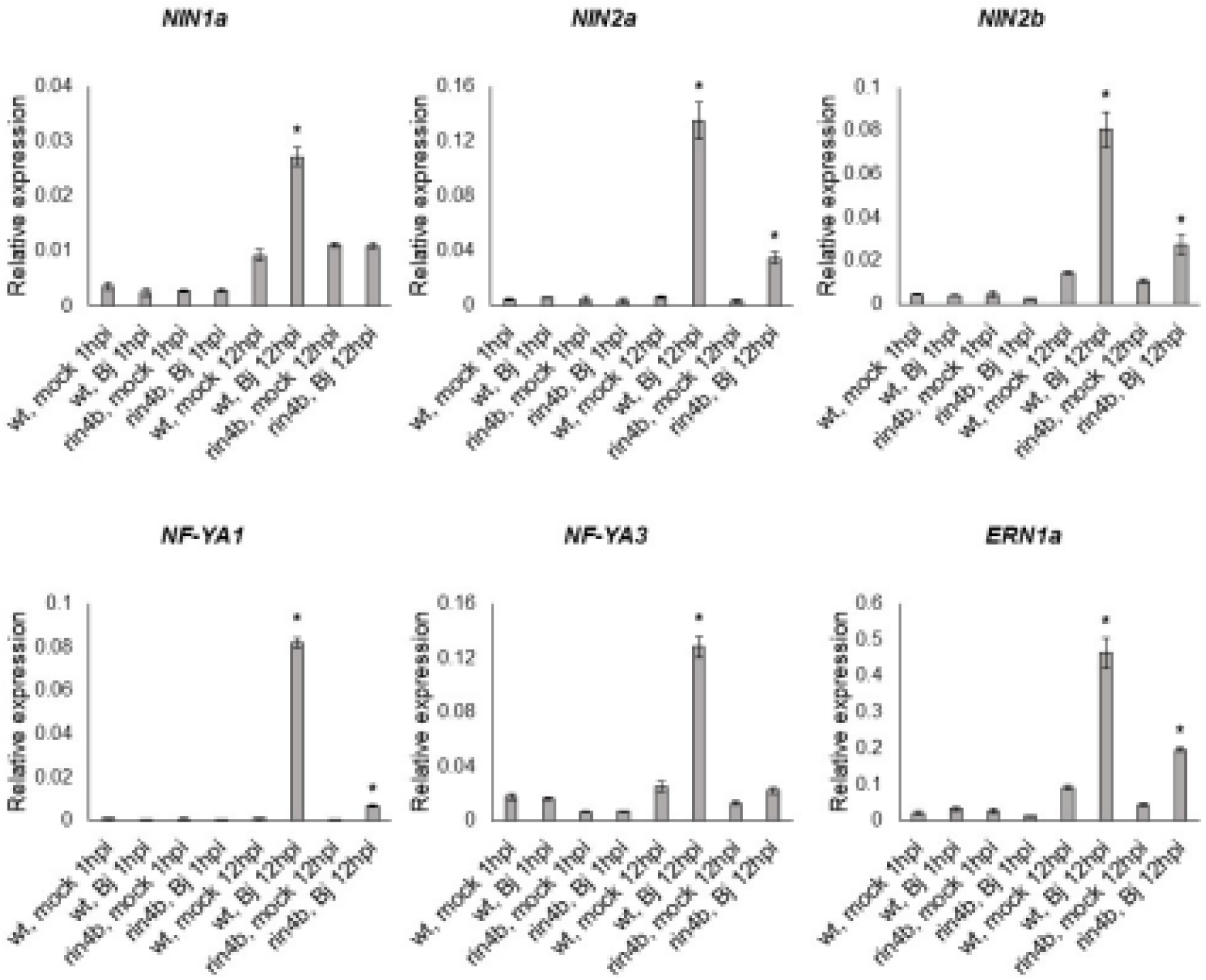
Symbiotic signaling is affected in the *rin4b*-CRISPR-Cas9 mutant. Expression of transcription factors (TF) involved in the symbiotic signaling pathway was investigated in the *rin4b* mutant background by qRT-PCR. Induction of the closest homolog of nodulation-specific transcription factor *NIN1a* can be observed in wild type (wt) and it does not change in *rin4b* in response to *B. japonicum* (Bj) 12 hpi. While two other homologs (*NIN2a* and *NIN2b*) respond to Bj in *rin4b,* their induction is significantly lower in comparison to Bert wt. Two soybean *NF-YA* TFs were tested as they are activated by NIN, therefore their lower expression in response to Bj in *rin4b* was expected. *ERN1* is another TF downstream of NIN. Its induction in *rin4b* in response to Bj is significantly lower than the response in Bert wt. Student’s t-test * p<0.05.

Since the expression of all investigated transcription factors was impaired in the *rin4b* mutant background in comparison to Bert wild-type plants at 12 hpi (Fig. 5), the data suggest that the absence of *RIN4b* negatively affects the symbiotic signaling pathway. While the expression of these marker genes was reduced in the *rin4b* mutant plants, some induction upon inoculation was still apparent. We attribute these findings to the functional redundancy between *RIN4a* and *RIN4b*.

As an invading organism, rhizobia also elicit plant immune responses (48, 49), although transient in the case of compatible interactions. These responses need to be amplified or suppressed depending on whether the host senses the bacteria as friend or foe. RIN4 is an immune regulator that is a key intersection between PTI, effector-mediated defense suppression and NTI (24). The work presented here also supports a role for RIN4 as a key determinant in symbiotic signaling.

In Arabidopsis, absence of RIN4 enhances PTI responses. Overexpression of AtRIN4 leads to PTI inhibition as no callose deposition was observed when plants were treated with flg22 (22). The function of AtRIN4 is controlled by the specific phosphorylation status of the protein. In the case of soybean nodulation, silencing of *rin4* expression, mutagenesis (i.e., *rin4b*-CRISPR-Cas9 mutant) or disruption of S143 phosphorylation resulted in a significant reduction in nodulation. Absence of AtRIN4 leads to increased PTI responses and, therefore, it might be that GmRIN4b absence also causes enhanced PTI which might have contributed to significantly less nodules. However, this may be too simplistic an idea given the impact of *rin4* mutagenesis on nodule-related gene expression, as well as the lack of knowledge of other likely partners (e.g., interacting R proteins) that might also be playing a role. Compared to leaves there is also a paucity of data as to how plant roots (51), especially with regard to soybean, respond to pathogen infection and elicitation, as well as any functional role for RIN4.

While no redundancy was shown between RIN4a and RIN4b in plant-pathogenic interactions, our data suggest functional redundancy between the two isoforms in the symbiosis, pointing toward a likely scenario when immune responses in the aerial part differ from immune responses in the root. Several defense-responsive genes which are well described in leaf immune responses could not be confirmed to respond in a similar manner in root (51). It was also shown that plant immune responses in roots are compartmentalized and specialized (51). Clearly, there remains much to be discovered as to how RIN4, as well as other components of both the pathogen response and symbiotic response pathways, converge to ultimately distinguish friend from foe in the root.

In soybean, an effector-mediated hijacking of the symbiotic signaling revealed nodule formation in the absence of the NFR1 receptor. NF-deficient *Bradyrhizobium elkanii* was able to elicit nodule formation on *nfr1* mutant plants, whereas NF-deficient T3SS (Type III secretion system is required for effector proteins delivery) double mutant was not able to induce nodule formation (52), suggesting that effector protein(s) are required for successful nodulation in soybean. Taking into consideration that rhizobial effector(s) were able to mediate nodule formation and AtRIN4 up-regulated phosphorylation is a target for an effector protein, our proposed model of RIN4 function in the symbiosis is the following: GmRIN4^S143^ is phosphorylated by SymRK, a receptor whose structure is similar to known pattern recognition receptors (PRR) (though no ligand was shown for SymRK so far). Putative effector proteins injected by compatible rhizobium recognize phosphorylated S143, ensuring downstream symbiotic signaling perhaps via effector-mediated interaction of RIN4 to modulate basal immune response. Another likely scenario would be phosphorylation mediated interaction of RIN4a and RIN4b proteins with other downstream proteins which would require further investigation into protein complex formation upon rhizobial infection.

Taken together, our results indicate that successful development of the root nodule symbiosis requires cross-talk between NF-triggered symbiotic signaling and plant immune signaling mediated by RIN4. Based on data presented here, all this likely happening upstream of the transcriptional regulator CYCLOPS, a further piece of evidence (in addition that GmRIN4a/b phosphorylation was detected only in root hairs, where the infection takes place) that involvement of soybean RIN4 in nodulation is required early on in the signalling pathway. Furthermore, it is likely that soybean RIN4 phosphorylation by SYMRK at S143 modulates immune responses to enable symbiotic interaction with rhizobia.

Our data present important findings on the way to unraveling how legume plants prioritize beneficial interactions and suppressing immunity-related signals to enable symbiotic associations.

## Material and Methods

A detailed description of the methods used in this study can be found in SI Appendix, Supplementary Materials and Methods.

### Seedling growth, treatment and total root, root hair and stripped root collection

In order to obtain root hairs and stripped roots, and total roots for protein and RNA extraction, *Glycine max* cv. Williams 82, and *rin4b*-CRISPR-Cas9 in Bert background M4 and M5 seeds, and Bert wild-type seeds were used. Seeds were surface sterilized with 20% household bleach, left in 20% bleach for 10 min, then washed three times with sterile diH2O. Afterwards, the seeds were incubated for 10 min in 0.05% HCl and subsequently washed four times with sterile diH2O before sowing onto 1% water-agar plates (20 cm in diameter glass Petri dish, MG Scientific, WI, USA). Williams 82 seeds (used for root hair experiments) were germinated in a growth chamber (Conviron Growth Chamber PGR15) with 90% humidity at 28°C in the dark and three days old seedlings were used for treatment. *rin4b* and Bert wild-type seeds were germinated at room temperature (21-23°C) in the dark. Seedlings were spray-inoculated with an OD600 ∼ 0.2 of *B. japonicum* USDA110 wild type, and sterile water sprayed as mock. Root hairs were harvested 1 hpi by flash-freezing in liquid nitrogen, followed by stirring for 15 min, which shears the root hairs from the roots. Roots hairs were then isolated by filtration and both root hairs and stripped roots (i.e., roots with root hairs removed) stored separately at -80°C until further processing. As for *rin4b* and Bert wild-type plants, roots were inoculated 4 days post sowing with mock and *B. japonicum* USDA 110 with an OD600 ∼ 0.2, and entire roots harvested 1 hpi and 12 hpi, and stored at -80°C until further processing.

### Protein purification from soybean root hairs, roots and transgenic roots

Soybean roots were ground in liquid nitrogen using mortar and pestle. Ground tissue was transferred to an Eppendorf tube and 750 μl of extraction buffer (0.9 M Sucrose, 100 mM Tris HCl, pH 8.0; 10 mM EDTA, 0.4% ß-mercaptoethanol, 0.2% Triton-X 100, Plant protease inhibitor, P9599 and phosphatase inhibitor, P0044 from Sigma) was added. Samples were vortexed and incubated on ice for 10 min. Equal volume of Tris buffered phenol was added, samples were vortexed and incubated for at least 1 h at 4°C rotating. Samples were centrifuged for 6 min at 13000 rcf at 4°C. Upper phase was transferred to a new tube and one ml ice-cold precipitation buffer (0.1 M Ammonium acetate dissolved in high quality methanol) was added and proteins were precipitated overnight at -20°C. Next day, samples were centrifuged for 10 min at 3500 rcf at 4°C. Pellet was resuspended in 1 ml ice-cold precipitation buffer, vortexed and centrifuged (10 min, 3500 rcf, 4°C). This step was repeated. Afterwards, the pellet was washed with 80% ice-cold acetone, vortexed and centrifuged (10 min, 3500 rcf, 4°C), this step was repeated twice. The pellet was resuspended in 70% EtOH, vortexed, centrifuged and air dried. Proteins were resuspended in 8 M Urea (solubilized in 50mM Tris-HCl, pH 8.0) and protein concentration was measured using Pierce 660 nm Protein Assay (Thermo Scientific, Rockford, IL, USA) and used for SDS-PAGE and subsequent Immuno-blotting analysis.

### SDS-PAGE and Immuno-blot analysis

10% or 12% Tris-PAGE gel was prepared without SDS (0.375 M Tris, pH 8.8; 10-12% Bis Acrylamide (40%, 29:1), 0.1% APS, TEMED) to separate proteins and for subsequent Western blot assay. Proteins were transferred onto a PVDF membrane at 4°C over-night at 30 V. After transfer, the membrane was incubated in 1% BSA (Gold Biotechnology, St. Louis, MO, USA) in TBS-T (with 0.1% Tween-20 from Fisher Scientific) or 5% milk (fat-free skim milk, SACO Foods, Middleton, WI, USA) in TBS-T. Custom RIN4 antibody was generated by AnaSpec (Fremont, CA, USA) and used at 1:5000 dilution in 5% Milk. Custom RIN4 phosphorylation site specific peptide antibody, α-pS143 (GRDPSPQWEPKNSYD) were generated by GenScript (Piscataway, NJ, USA) and used at 1:4000 dilution in 1% BSA to detect proteins isolated from soybean root hair. Secondary rabbit HRP conjugated antibody was used in 1:10000 dilution and obtained from Jackson ImmunoResearch (West Grove, PA, USA).

HA-tagged RIN4a and RIN4b expressed in soybean transgenic roots were detected using monoclonal HA-antibody (Roche Diagnostics, Germany) with or without HRP-conjugate and used in 1:2500 dilution. 1:5000 dilution was applied to detect HA-tagged RIN4a/b expressed in protoplasts. In case of non-conjugated HA antibody, secondary rat antibody was used in 1:10000 dilution in 5% milk. Signal was visualized using 1:8 Femto:Pico Pierce SuperSignal chemiluminescence substrate (Thermo Fisher Scientific, USA). GFP expressed in transgenic roots was detected using anti GFP antibody (Invitrogen, USA) at 1:5000 dilution in 5% milk. To detect RIN4(a/b) S143 phosphorylation in protoplasts, membrane(s) were blocked in a 2% BSA (dissolved in TBS-T) solution, and were incubated with α-pS143 at 1:3000 in 2% BSA solution with subsequent incubation with secondary rabbit HRP conjugated antibody at 1:15000 dilution. Pictures were taken using a UVP Camera (BioSpectrum 815 Imaging System; Upland, CA, USA) system, or x-ray film was exposed by the blot and photographically developed. Images were inverted and brightness and contrast were adjusted during figure preparation.

### Targeted Multiple Reaction Monitoring Mass Spectrometry (MRM-MS)

Costume-made stable-isotope-labeled AQUA peptides carrying the S143 and T173 phosphorylation sites (RIN4a and RIN4b are identical, IS Appendix, Table S3) were generated by Sigma-Aldrich (The Woodlands, TX, USA). The peptides were resuspended in 50% Acetonitrile (ACN) to generate 10 pmol/µl stock solutions stored at -80°C. Working solutions of the peptides were diluted in 5% ACN with 0.1% formic acid (FA). After 30 min *in vitro* kinase assay (described above), the samples were spun down and a mixture of heavy-labeled S143, pS143 (phosphorylated version of S143 peptide), T173 and pT173 were added (100 fmol of S143 and 50 fmol of T173) to the samples in 20 µl reaction volume prior to digestion. In solution trypsin digestion was performed overnight at 37°C as follows: 2.5 µl of reduction solution (100 mM Ammonium bicarbonate/ABC with 100 mM DTT) was added, and samples were incubated at 37°C for 30 min; 2 µl alkylation solution (0.5 M Iodoacetamide dissolved in 100 mM ABC) was added to the samples and samples were incubated up to 1 hour at room temperature in the dark. Samples were neutralized by 80 ul 10 mM DTT in 10 mM ABC solution, before 10 µl sequencing grade modified Trypsin (100 ng/µl) from Promega (Madison, WI, USA) dissolved in 100 mM ABC was added. After overnight trypsin digestion, samples were centrifuged for 2 min at 13 000 rcf, frozen in liquid nitrogen and lyophilized. Samples were dissolved in 0.1% FA. To quantify phosphorylation, we used Thermo-Scientific Quantiva triple Quadrupole mass spectrometer coupled to an Eksigent 1D plus (SCIEX) Nano-LC (liquid chromatography) instrument. 20 cm long column of 75 µm in diameter filled with HxSIL-C18 (particle size 5 µm, Hamilton Company, Reno, NV, USA) was used for sample separation over a 25 min gradient run. 10 μl sample per injection was used in three technical replicates for each biological replicate. MS-MRM (in positive ion mode) was run over 25 min acquisition time at 3 mTorr CID gas pressure, defined collision energy (V) and cycle time (1 sec) for at least three transitions for each peptide (IS Appendix, Table S2). MS RAW data files were processed using Skyline software (MacCoss Laboratory Software, University of Washington, Seattle, WA, USA; https://skyline.ms/project/home/begin.view<u>?</u>) to obtain the area under curves for integrated LC-SRM peaks. Integrated peaks were manually inspected to ensure all quantified transitions had overlapping retention times. Native peptide abundance is expressed as the ratio of endogenous peptide to labeled standard peptide expressed in percentage. MS-SRM was performed at the Gehrke Proteomics Center of the University of Missouri.

## Acknowledgements

We thank Dr. Brian Mooney (Associate Director of MU Gehrke Proteomics Center) for his patience and help with MS-SRM and comments on the respective part of the manuscript. The authors are grateful to Yer Xiong for transforming the *RIN4b*-CRISPR-Cas9 construct into soybean. (University of Minnesota, St. Paul, MN, USA). We also thank Dr. Jean-Michel Ane (University of Wisconsin, Madison, WI, USA) and Dr. Katharina Pawlowski (Stockholm University, Sweden) for their help with RIN4 sequences from FaFaCuRo species published in Griesmann *et al*., 2018; Salgado *et al*., 2018 and Leebens-Mack *et al*., 2019. We also want to express our appreciation for the help of MU Molecular Cytology Core with Confocal Laser Scanning Microscopy.

This research was supported by the National Science Foundation (NSF) Plant Genome Program under award number IOS-1734145 and by an FY18 Mini Research Grant to KT from MU Gehrke Proteomics Center.

## Materials and Methods

### Sequence alignment and phylogenetic analysis of RIN4 protein homologs

Putative RIN4 homologs were defined with at least one NOI domain (PFAM database ID: PF05627 called AvrRpt-cleavage family) in protein sequences. Protein sequences were downloaded from the Phytozome database (http://phytozome.jgi.doe.gov/), (http://www.kazusa.or.jp/lotus/), NCBI database (1, https://www.ncbi.nlm.nih.gov/), PeanutBase (https://www.peanutbase.org/home), *Lotus japonicus* database (https://www.kazusa.or.jp/lotus/) and the UniProt protein database (https://www.uniprot.org/), from GigaDB (34, https://www.re3data.org/repository/r3d100010478) and from 1kp project (2, https://db.cngb.org/onekp/). All predicted RIN4 homologs were confirmed with at least one NOI domain using Simple Modular Architecture Research Tool (SMART, http://smart.emblheidelberg.de/) and manual check. A full-length alignment of all putative RIN4s were made using the MAFFT server tool (http://mafft.cbrc.jp/alignment/server/) with iterative refinement methods (E-INS-i), multiple conserved domain alignment option, scoring matrix BLOSUM62, default gap opening penalty 1.53.

The alignments were used to generate phylogenetic trees. The phylogenetic tree in the Fig. 1A was generated using the Neighbor Joining (NJ) method (3) with the JTT substitution model (4) and 1000 bootstrap resampling value. The phylogenetic tree in the Fig. S1 was generated using the Average linkage (UPGMA) method and the JTT substitution model. Phylogenetic tree was visualized using the FigTree tool.

### Composite plant generation and nodulation assay

In order to generate soybean composite plants expressing the respective constructs, Agrobacterium-mediated hairy root transformation was performed as described in Tóth *et al*, 2016 (5) with the following modifications. Hairy root induction was initiated by poking the plants with a needle tip (BD PrecisionGlide Needle, 23G x 1½ TW IM/0.6mmx40mm, sterile, Franklin Lakes, NJ, USA) carrying the respective agrobacterium. Before planting the composite plants into sterile potting mix (3:1 vermiculite: perlite mix; Hummert International, Earth City, MO, USA), Agrobacterium induced roots were subjected to fluorescence microscopy. The vectors used in this study contain a constitutive GFP marker in order to identify transgenic roots. Non-transgenic roots were removed before planting. 24-48 hours after planting, plants were inoculated with B. *japonicum* USDA 110 wild type at an OD600 ∼ 0.05. Composite plants were grown in a walk-in growth chamber (Conviron GR 64; non-controlled humidity; 16 h/light/26°C and 8 h/dark/24°C). The nodulation phenotype was observed at 5 wpi. Leica M205 FA stereomicroscope was used to take pictures of the nodulated transgenic roots at 8.0 x magnification.

In order to determine the nodulation phenotype of *rin4b* mutant plants, seeds (M4) of *rin4b* homozygous CRISPR-Cas9 lines and non-edited Bert plants, with native *RIN4b* (originating from plants that went through transformation but segregated as non-transgenic and non-edited), were surface sterilized and seedlings transferred to perlite-vermiculite potting mix and were grown in controlled walk-in growth chamber (Conviron GR 64; non-controlled humidity non-controlled humidity; 16 h/light/23°C and 8 h/dark/21°C). Plants were inoculated with B. *japonicum* USDA 110 wild type at an OD600 ∼ 0.05 and subjected to phenotyping 5 wpi. A Panasonic (Lumix) camera was used to take pictures of Bert wild-type and *rin4b* mutant nodulated roots.

### CRISPR-Cas9 edited *RIN4b* (Glyma16G090700) mutant generation

The transformation vector was constructed using a modified version of the *Glycine max* codon optimized Cas9 plasmid from Michno *et al*. (6, Addgene plasmid # 59184). The pBlu guide RNA shuttle vector used was identical to the one described in Michno et al. (6) (addgene #59188). *RIN4b* target was selected using Stupar lab’s CRISPR design tool (http://stuparcrispr.cfans.umn.edu/CRISPR/). Guide RNA target oligos were designed based on the program then synthesized and annealed in a 10X PCR buffer at 50 °C for six hours. The pBlu/gRNA shuttle vector was digested using BbsI (New England Biolabs # R0539S) following manufacturer’s guidelines. Digested product was run on an agarose gel and extracted using Qiagen gel extraction kit. The target oligos and shuttle vector were ligated using T4 ligase (New England Biolabs # M0202S). The ligation product was transformed into DH5 alpha (Life technologies) competent cells. The pBlu vector containing the target oligos and the destination vector were both digested with EcoRI (New England Biolabs #R0101S). The gRNA cassette and digested destination vector were ligated using T4 ligase, and transformed into DH5 alpha competent cells. *RIN4b* CRISPR-Cas9 construct was delivered into Bert-MN-01 background using K599 Agrobacterium (disarmed strain 18rv12). Methods for delivery and growth of whole-plant transformants were performed as previously described

(7). T0 plants and subsequent progeny were tested for the presence of the CRISPR-Cas9 transgene using PCR with transgene-specific primers followed by agarose gel electrophoresis. Mutations in transformed plants were screened as previously described using cleaved amplified polymorphic sequences (8) and/or PCR heteroduplex (9, 10) analysis. The mutated alleles in specific plants were confirmed by Sanger sequencing of PCR amplicons. Plants harboring deletions and no transgene were selected for further analyses.

### Cloning

*GmRIN4a* (Glyma.03G084000.1) and *GmRIN4b* (Glyma.16G090700.1), GmSymRKß (Glyma.09G202300.1), GmNFR1α (Glyma.02G270800.1) CDS were cloned via Gateway BP reaction into pDONR/Zeo entry vector. Sequence accuracy was confirmed by Sanger sequencing performed by MU DNA Core facility. These entry clones were used to fuse the haemagglutinin (HA) epitope onto the N-terminus of RIN4a and RIN4b and to introduce point-mutations by site-directed mutagenesis. HA-epitope tagged entry clones were used for subsequent cloning into a modified, gateway compatible pCAMBIA vector (11) for ectopic expression in soybean composite plants. 120-150 bp of *RIN4a* and *RIN4b* transcripts in the 3’UTR regions were cloned into the pDONR/Zeo entry vector. These clones were used for subsequent cloning into a modified gateway compatible pCAMBIA vector for RNAi-mediated gene silencing (12). Both modified pCambia vectors contain a GFP marker for transgenic root identification. pDONR/Zeo: RIN4a/b and pDONR/Zeo: SymRKß/NFR1α entry clones were used to create constructs for BiFC, Split-Luciferase, as well as protoplast (protein expression and phosphorylation) experiments.

### Bimolecular Fluorescence Complementation Assay

pDONR/Zeo:*RIN4a* and *RIN4b* were used to create N-terminally fused split YFP constructs in the pAM-PAT vector series (13), and translational fusion was created with the N-terminal as well as C-terminal half of the split YFP fluorophore. NFR1α, SymRKßΔMLD and P2K1 were fused C-terminally to the C-terminal YFP domain. *Nicotiana benthamiana* leaves were infiltrated with *Agrobacterium tumefaciens* GV3101 (grown in LB media supplemented with 50 μg/ml Carbenicillin and 20 μg/ml Gentamicin) carrying the respective constructs and p19 (100 μg/ml Kanamycin and 25 μg/ml Rifampicin) silencing suppressor.

Microscopy was performed 40-46 hours post infiltration. A LEICA SP8 confocal laser scanning microscope with a tunable white light laser (WLL) was used to visualize YFP fluorescence generated upon interaction between the co-expressed proteins as a result of split YFP halves reconstitution. Plasma membrane (PM) was labeled with FM4-64 (Invitrogen, USA) PM dye. YFP was excited at 514 nm and the emission was detected at a 525-575 nm bandwidth. FM4-64 was excited at 510 nm and dye’s emission was detected using a 700-780 nm bandwidth. Images of two fluorescence channels were acquired sequentially with a 40x/1.1NA water immersion objective and an additional zoom factor 3. The pixel size of images was set to 95 nm. Brightness and contrast of the images was adjusted in PowerPoint. CLSM was done at the Molecular Cytology Core of the University of Missouri.

### Split-Luciferase Complementation Assay

RIN4a/b as well as NFR1α, SymRKßΔMLD and P2K1 were C-terminally fused to split halves of Luciferase in a Split-Luciferase vector system (pCAMBIA-GW-Nluc and pCAMBIA-GW Cluc) and the following constructs were generated via LR reaction: RIN4a/b:Cluc; NFR1α:Nluc, SymRKßΔMLD:Nluc, P2K1:Nluc, and transformed into *Agrobacterium tumefaciens* GV3101 strain. Leaves of 3-weeks old tobacco plants were co-infiltrated (Infiltration buffer: half Murashige and Skoog liquid media and 150 μM Acetosyringone) with agrobacterium carrying the respective constructs at OD600 ∼0.55 together with agrobacterium strain carrying p19 silencing suppressor. Protein-protein interaction was observed 2 or 3 dpi, via LUC activity, when Luciferin buffer [100 mM Tris-HCl pH 7.8, 5 mM MgCl2, 0.15 mM ATP (Sigma), 5 mM D-Luciferin (GoldBio), and 0.01% Silwet-L-77] was sprayed onto the leaves, incubated in dark for 7 min (to decrease autofluorescence originating from chloroplasts) and luminescence was captured by a CCD camera (Photek 216; Photek, Ltd.).

### Recombinant protein expression and *in vitro* kinase assay

GmRIN4a, GmRIN4b, GmRIN4c (Glyma.18G166800.1) and GmRIN4d (Glyma.08G349500.1) were fused C-terminally with a His-epitope in the pET22b vector (Novagen). The intracellular, kinase domains of GmSymRKβ (537 aa - 919 aa) and NFR1α (294 aa - 599 aa) were N-terminally tagged with GST-epitope in the pGEX-5X-1 vector (GE Healthcare, Pittsburgh, PA, USA). Rosetta^TM^ (DE3) bacterial cells (Novagen/Sigma-Aldrich, Saint Louis, MO, USA) carrying the respective constructs were grown in LB medium (with respective antibiotic) and cell cultures were induced with 0.5 mM isopropyl β-D-1- thiogalactopyranoside (IPTG) after reaching an OD600 absorbance of 0.5 and incubated at 28°C for an additional 4 hours. Bacterial cells were collected at 4,000 rpm for 10 min and His and GST-tagged proteins were purified by TALON Metal Affinity Resin (Clontech, Takara Bio, USA) and Glutathione Resin (GenScript, Piscataway, NJ, USA) following the manufacturer’s protocol, respectively. For the *in vitro* kinase assay, 2 μg of purified GST tagged protein kinases were incubated with 2 μg His-tagged GmRIN4 proteins as substrate in a 20 μl reaction buffer containing 20 mM Hepes-KOH (pH 7.4), 5 mM MgCl2, 1 mM DTT, 2 mM ATP, and w/wo 0.2 μl radioactive [γ-32P] ATP for 60 min at 30°C. In the case of radioactive detection, 5 μl of 5× SDS loading buffer was added to the reaction, and samples were boiled for 5 min. The proteins were separated by electrophoresis in 12% SDS-PAGE gels, followed by autoradiography for 12 h. The proteins were visualized by staining the gel with Coomassie Brilliant Blue (CBB) and auto-radiographed using a Typhoon FLA 9000 phosphorimager (GE Healthcare). Myelin Basic Protein (MBP) and GST were used as controls.

### Protein expression and *in planta* phosphorylation in *Arabidopsis* protoplasts

To assess RIN4a and RIN4b phosphorylation *in planta*, the proteins were fused to HA-epitope in a pUGW14 vector driven by 35S promoter and co-expressed with SYMRKΔMLD (wild type and kinase-dead versions) fused to HA-epitope in *Arabidopsis* leaf protoplasts. Protoplast isolation, transfection and protein extraction from protoplast was performed as described in Cho *et al*., 2022 (14).

### RNA extraction and qRT-PCR experiment

RNA was extracted from entire soybean roots, root hairs and stripped roots, and from transgenic roots using Trizol (Sigma, St. Louis, MO, USA). Tissue was ground in liquid nitrogen, 1 ml 1.5 ml Trizol reagent was added to the mortar and samples were transferred into an Eppendorf tube as liquid. These extracts were incubated on ice for 10 min and then centrifuged at 13000 rcf for 10 min at 4°C. The supernatant was transferred into a new tube and 200 μl chloroform was added per 1 ml supernatant, vortexed and centrifuged for 20 min at 4°C. The upper phase was carefully transferred into a new tube and the half volume of cold ethanol was added. From this step forward, the samples were transferred onto a Qiagen column, using Qiagen RNeasy extraction kit (Qiagen, Hilden, Germany), following the manufacturer’s protocol. The quality of the RNA was checked by agarose-gel electrophoresis and samples were DNAse treated using Ambion Turbo DNAse (Invitrogen by Fisher Scientific, Vilnius, Lithuania) following the manufacturer’s recommendation. cDNA synthesis was performed using 2 μg RNA, oligo dT primer, and Promega MLV Reverse transcriptase (RT) kit (Madison, WI, USA), a negative control without RT was included in cDNA-synthesis. For qRT-PCR, cDNA was diluted five times and Applied Biosystems SYBR Green (Thermo Fisher Scientific, USA) was used to perform quantitative RT-PCR. For data analysis, Rn values were extracted from ABI 7500 Real Time PCR machine and LinReg software (https://www.gene-quantification.de/LinRegPCR_help_manual_v11.0.pdf) was used to determine baseline and Cq values. Data were extracted into Excel file and *Cons6* (F-box protein encoding gene) and/or *Cons7* (Insulin-degrading enzyme, Metalloprotease) was used as a reference gene (15) to normalize Cq values. ΔCt method (16) was used to evaluate the data and determine relative expression. qPCR primers were designed using Primer3 PCR primer design tool (17). Primers were designed based on the Williams 82 reference genome. qRT-PCR analysis was performed on cDNA derived from Williams 82 root hairs and stripped roots, and cDNA derived from total root of Bert cultivar and *rin4b* mutant in Bert background.

---

**Fig. S1.**
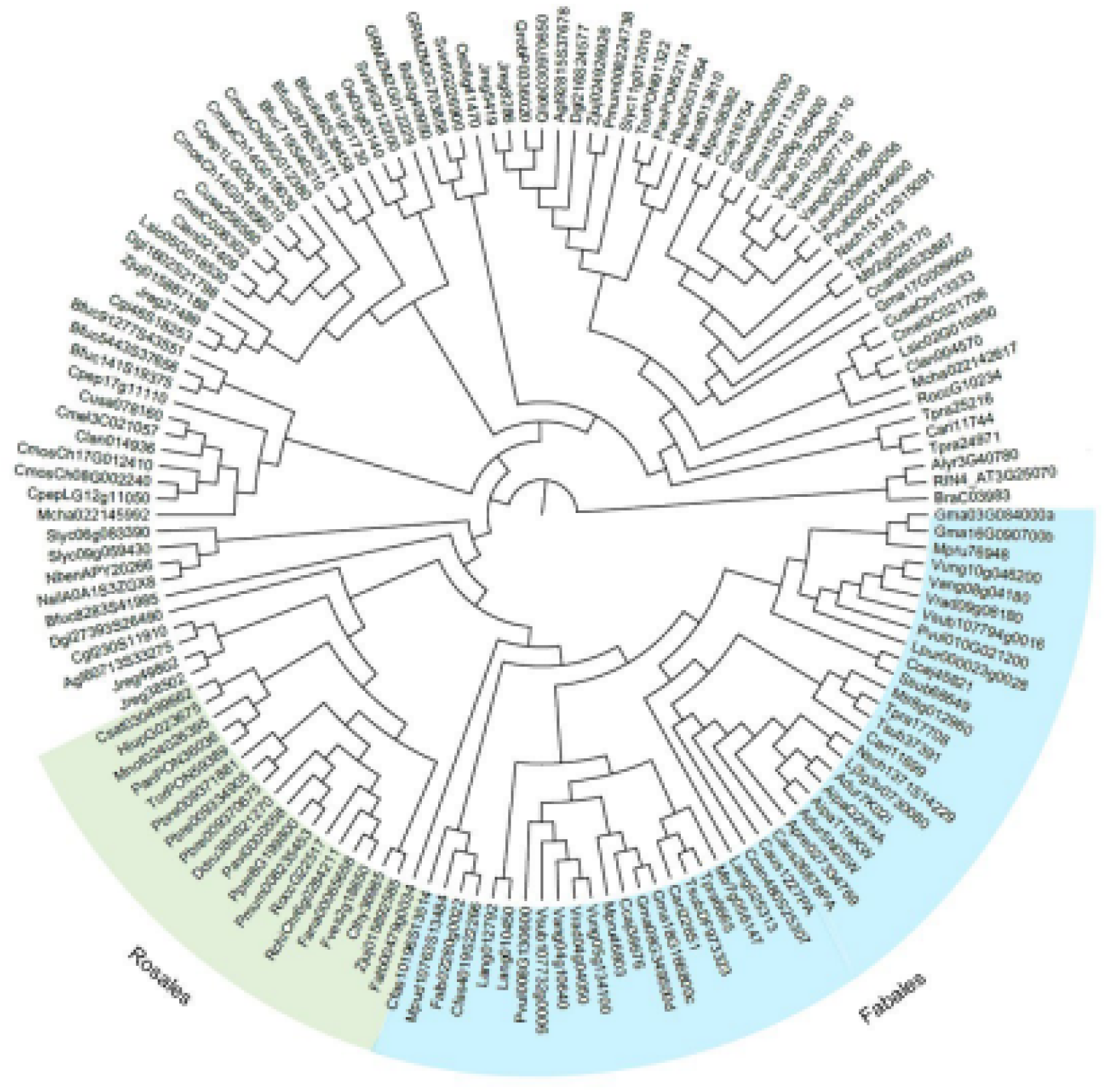
Phylogenetic tree of RIN4 proteins shows a nodulation-specific subclade. Phylogenetic tree of 149 closest *RIN4* homologs from 66 species, from non-legumes such as *Arabidopsis thaliana*, *A. lyrata, Brassica rapa*, *Brachypodium distachyon*, *Setaria viridis, Zea mays*, *Oryza sativa, Nicotiana benthamiana, N. silvestris, Solanum lycopersicum.* and from the phylogenetic FaFaCuRo clade: Fabales (nodulating species: *Abrus precatorius, Arachis ipaensis*, *A. duranensis, Cajanus cajan, Chamaecrista fasciculata*, *Cicer arietinum, Faidherbia albida, Glycine max*, *Lablab purpureus, Lotus japonicus*, *Lupinus angustifolius*, *Medicago truncatula*, *Mimosa pudica*, *Mucuna pruriens, Phaseolus vulgaris*, *Spatholobus suberectus, Trifolium pratense*, *T. subterraneum, Vigna radiata*, *V. angularis*, *V. unguiculata, V. subterranean;* non-nodulating species: *Chastanospermum australe, Cercis canadensis*, *Nissolia schottii*), Rosales (*Parasponia andersonii* nodulating with rhizobia, its non-nodulating relative *Trema orientale*; *Dryas drummondii* nodulating with Frankia, and non-nodulating *Cannabis sativa, Ceanothus thyrsiflorus*, *Fragaria vesca, Fragaria x ananassa, Humulus lupulus, Morus notabilis, Prunus persica, P. mume, P. avium, Pyrus bretschneideri, Rubus occidentalis, Rosa chinensis, Ziziphus jujube*), Fagales (nodulating *Alnus glutinosa, Casuarina glauca* and non-nodulating *Junglans regia, Quercus robur, Q. loboa*), and Cucurbitales (nodulating *Datisca glomerata*, non-nodulating *Begonia fuchsioides, Citrullus lanatus, Cucumis sativus, C. pepo, C. melo, Lagenaria siceraria, Momordica charantia*) (37, 52). Two sub-subclades formed: the blue-highlighted contain all the legume RIN4 homologs, whereas the green contains RIN4 homologs from Rosales. The tree was build using Average linkage (UPGMA) method and JTT substitution model.

**Fig. S2.**
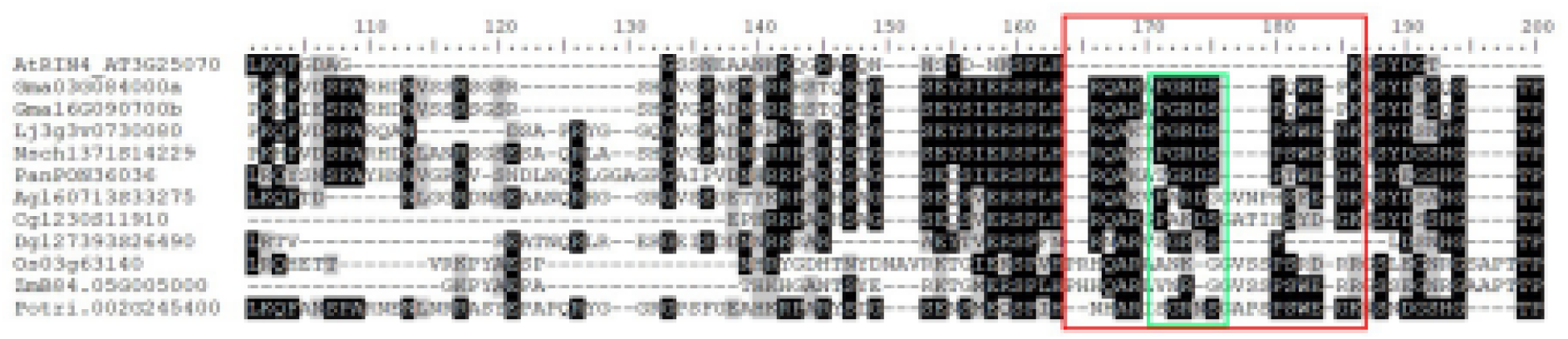
Alignment of representative RIN4 homologs from the FaFaCuRo clade. The FaFaCuRo clade contains species which are able to form symbiotic nitrogen fixation. Here, we show that the 15 amino acid novel-RIN4-motif (NRM) sequence (red box) and its “GRDSP” core motif (green box) – which is present in soybean (Gma03G084000/RIN4a, Gma16G090700/RIN4b), *L. japonicus* (Lj3g3v0730080), *N. schottii* (Nsch1371S14229) and Parasponia (PanPON36036) RIN4 proteins – is not conserved in nodulating species from Fagales (*Alnus glutinosa* – Agl160713S33275; *Casuarina glauca* – Cgl1230S11910) and Cucurbitales (*Datisca glomerata* – Dgl127393S26490). In addition, non-Fabales species: rice (*Oryza sativa –* Os03g63140), maize (*Zea mays* – ZmB84.05G005000) and poplar (*Populus trichocarpa –* Potri.002G245400) were included in the alignment.

**Fig. S3.**
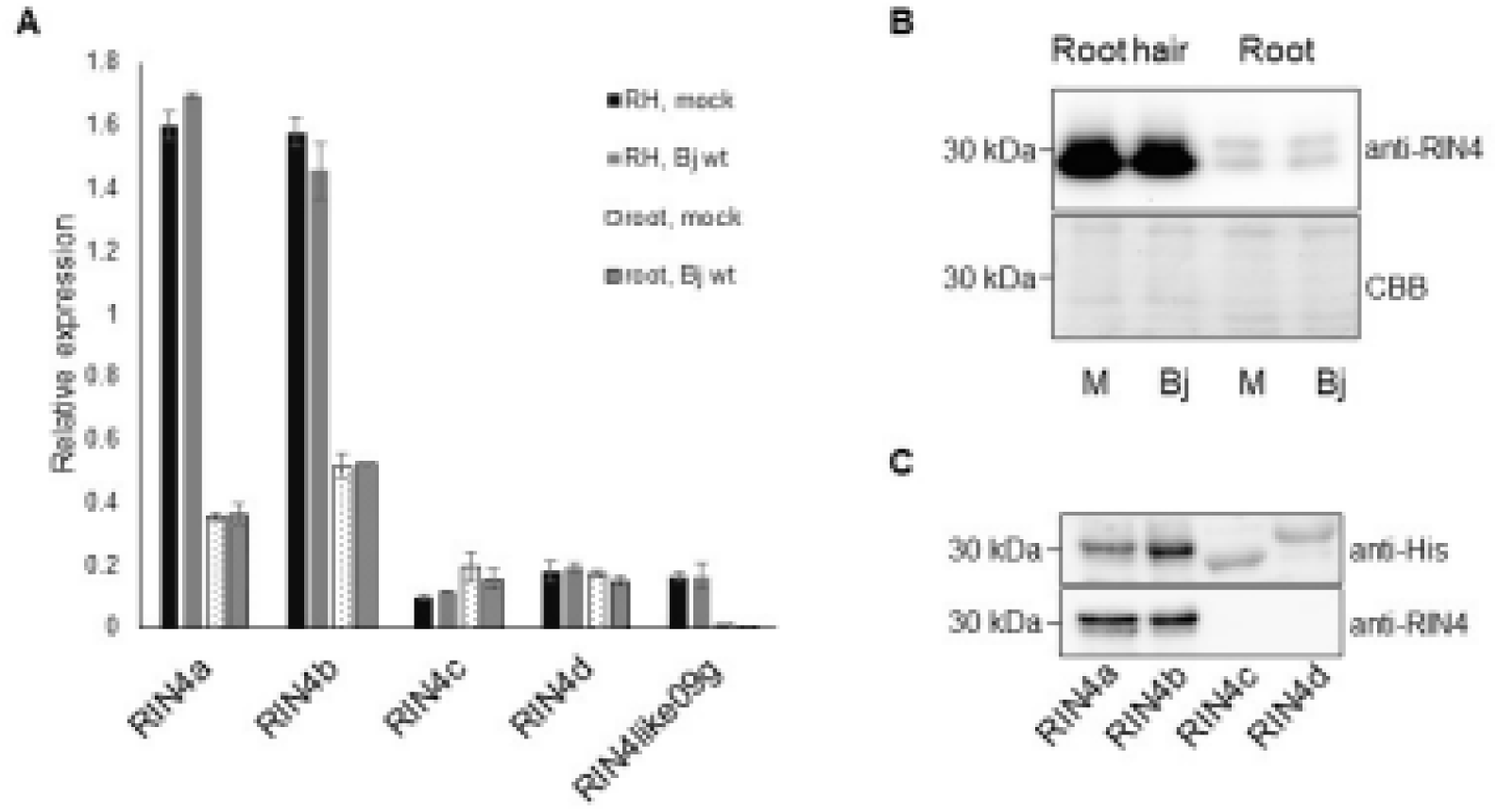
Soybean RIN4a and RIN4b are highly expressed in root hair and stripped root. (A) Relative gene expression levels of soybean *RIN4* homologs (*RIN4a-d*) and a *RIN4-like* gene (*Glyma09G008700*). *RIN4a* and *RIN4b* display higher expression levels than *RIN4c*, *RIN4d* and the *RIN4-like* gene in root hair (RH). All four have lower expression levels in roots. No difference could be observed between mock and rhizobial (Bj wt) treatment in root hairs or in roots. qRT-PCR analysis was done on 3 biological replicates, data show the mean of 2 technical replicates of 1 biological replicate. (B) Immuno-blot analysis, RIN4 proteins detected using RIN4-specific antibody, shows higher protein levels in root hairs than in roots (upper panel). No response was observed to treatment with *B. japonicum* (Bj) in comparison to mock treatment (M). Lower panel: Coomassie Brilliant Blue (CBB) staining of the same membrane showing equal loading. Immuno-blot analysis was performed on three biological replicates. (C) Custom-made anti-RIN4 antibody was tested on recombinantly expressed His-epitope tagged RIN4a, RIN4b, RIN4c and RIN4d proteins. Anti-RIN4 can recognize only RIN4a and RIN4b proteins. Upper panel: Immuno-blot showing RIN4a, b, c, d proteins detected using anti-His antibody. Bottom panel: Immuno-blot detecting RIN4 proteins using anti-RIN4 antibody.

**Fig. S4.**
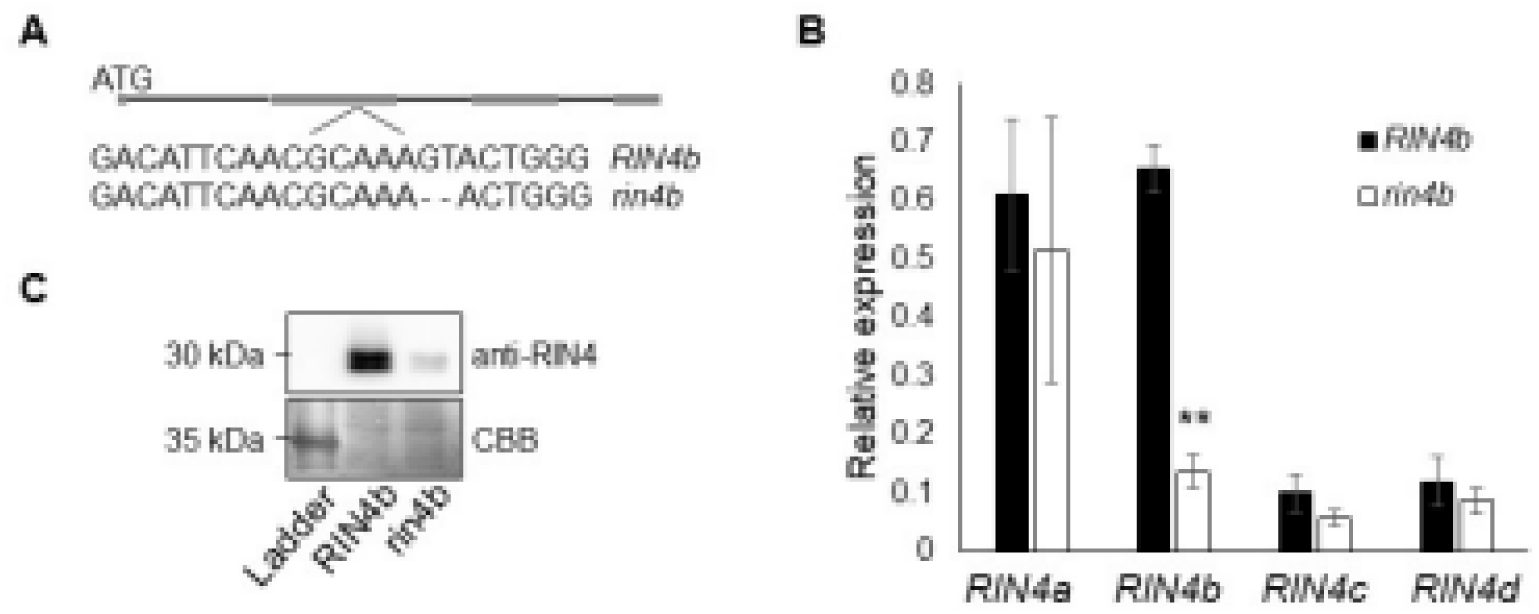
*Rin4b*-CRISPR-Cas9 deletion and reduced mRNA and protein level in *rin4b*-CRISPR-Cas9 mutant. (A) Two base pairs deletion was introduced in the second exon of *RIN4b* (in Bert background) using CRISPR-Cas9 technology which led to a pre-mature stop codon in *rin4b* mutant. (B) qRT-PCR analysis performed on roots shows that *RIN4b* mRNA levels were reduced in *rin4* mutant in comparison to wild-type roots (*RIN4b*), while *RIN4a*, *RIN4c* and *RIN4d* levels were not affected in the *rin4b* mutant background. Error bars represent standard error. Student t-test ** p<0.005. (C) RIN4 protein abundance was reduced in *rin4b*-CRISPR-Cas9 mutant roots (*rin4b*) in comparison to wild type roots (RIN4b). Immuno-blot detection was performed using anti-RIN4 antibody on total protein extracted from roots (upper panel). Lower panel: Membrane stained with Coomassie Brilliant Blue (CBB) to show loading.

**Fig. S5.**
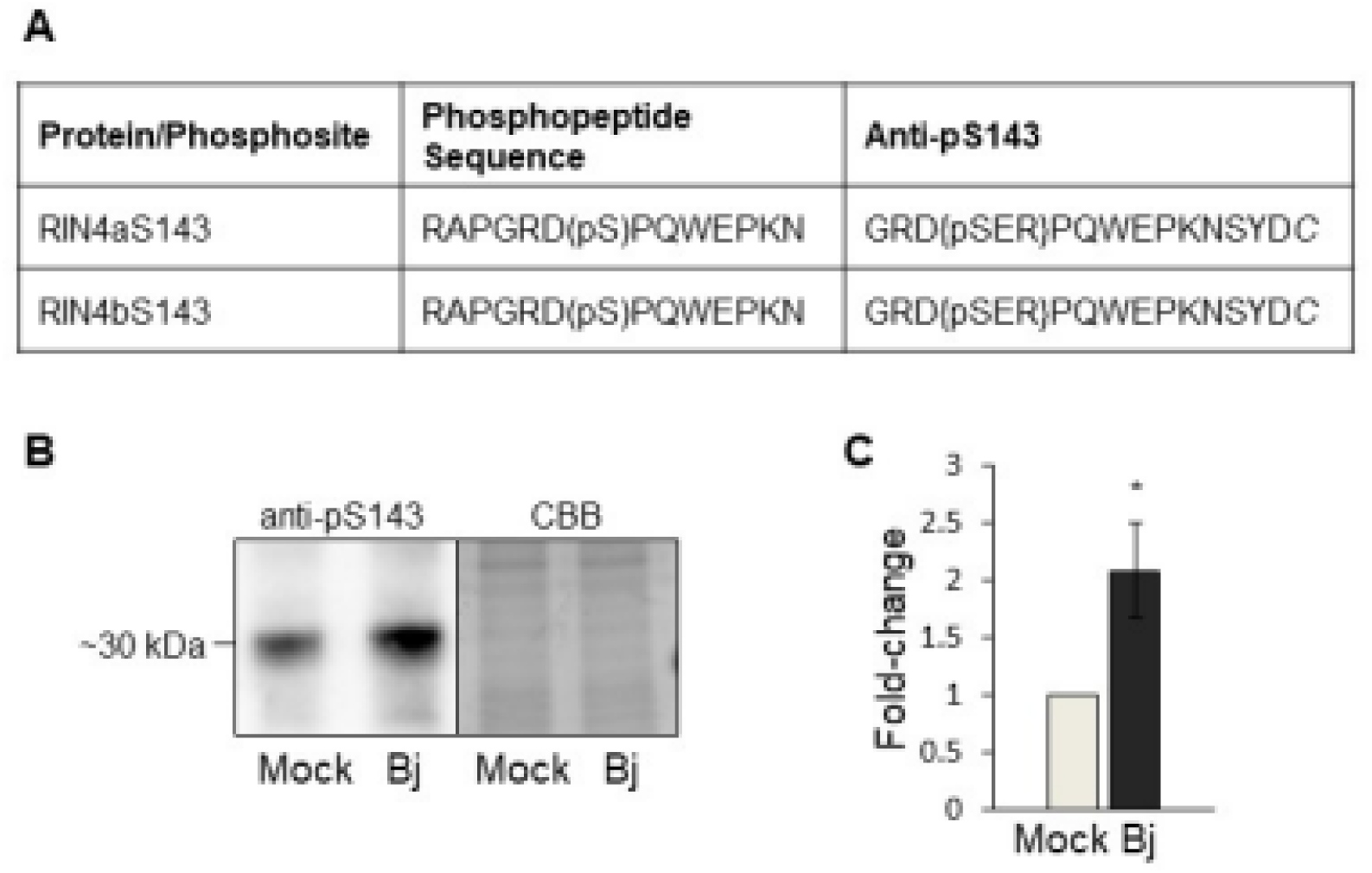
Nodulation-related S143 phosphorylation site is upregulated in response to rhizobium. (A) Table describing the sequence of the phosphopeptide identified in our previous study (Nguyen *et al*., 2012). The peptide sequence carrying the S143 phosphorylation sites is identical in RIN4a and RIN4b proteins. The right column shows the sequence of the anti-phosphopeptide against the sequence carrying the S143 phosphorylation site. A cysteine “*C*” is automatically added to a peptide during synthesis. (B) Phosphorylation of RIN4^S143^ is up-regulated in soybean root hairs in response to soybean symbiont *B. japonicum*. Left panel: Immune-blot detection using pS143 peptide antibody performed on root hairs treated with H2O (mock) and soybean symbiont *B. japonicum* (Bj), 1 hpi. Right panel: Coomassie Brilliant Blue (CBB) staining of protein gel previously run to determine equal loading. (C) Quantification of phosphorylation using ImageJ software. Error bar represents standard error. Experiment was repeated three times.

**Fig. S6.**
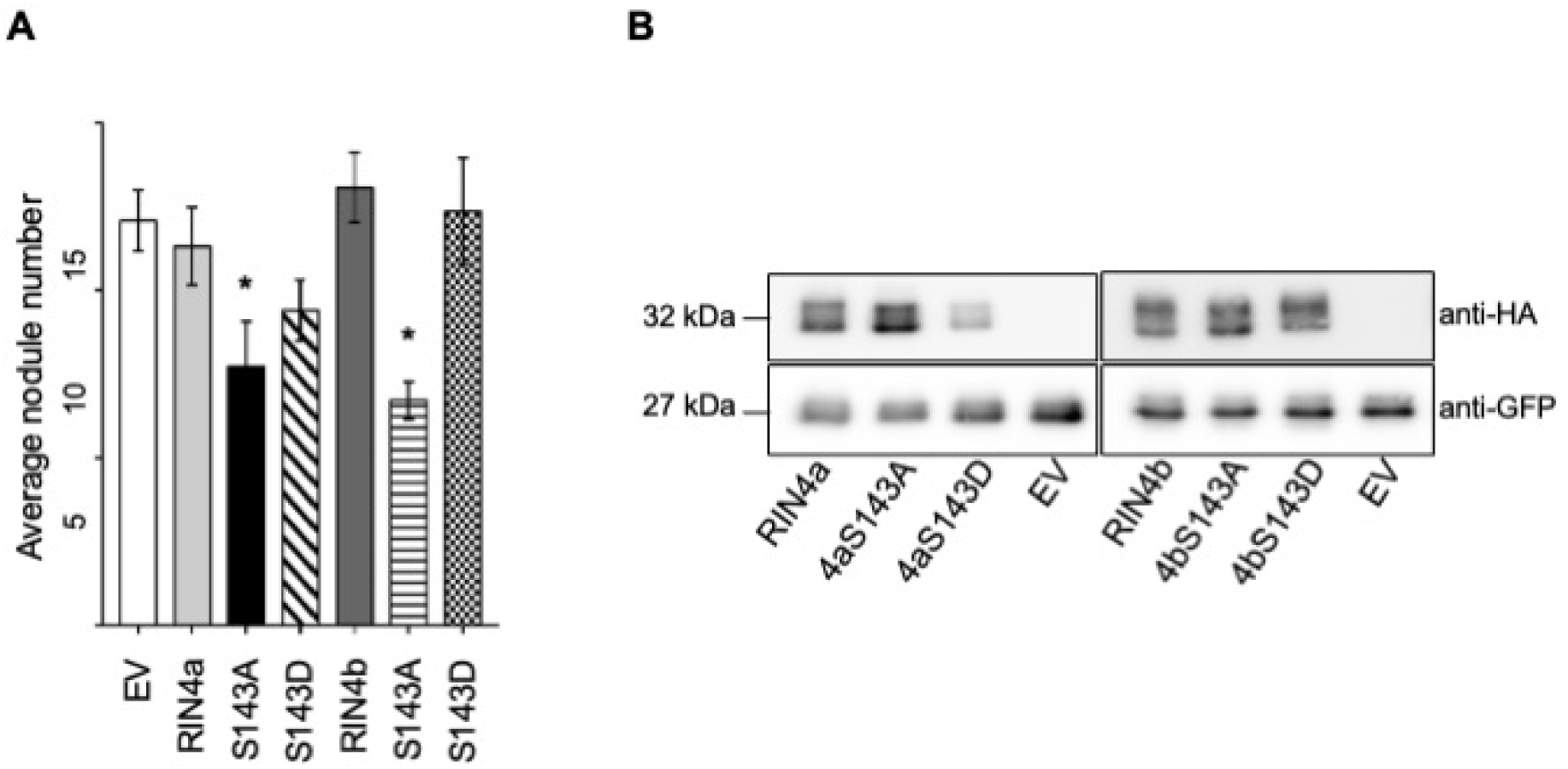
Novel S143 phosphorylation site contributes to symbiosis development. (A) Phosphor-minus (Ala; S143A) and phosphomimic (Asp; S413D) mutations were introduced into RIN4a and RIN4b to replace S143 residue, and the constructs were expressed in soybean transgenic roots. RIN4a^S143A^ and RIN4b^S143A^ displayed significantly reduced nodule numbers in comparison to transgenic roots carrying empty vector (EV) and the wild-type RIN4a and b protein. RIN4a^S143D^ and RIN4b^S143D^ did not have an effect on nodulation. Roots were phenotyped 5 wpi. Experiment was repeated three times with similar results. Error bars represent standard error. Student’s t-test * p<0.05. (B) Expression of HA-epitope tagged RIN4a and RIN4b and their phosphor-minus (S143A) and phosphor-mimic (S143D) versions in soybean transgenic roots. Immuno-blot analysis confirmed the expression of each version of RIN4a and b. Upper panel: HA antibody detecting RIN4a and b native and mutated proteins. Lower panel: detecting free GFP (27kDa) used as a marker to detect transgenic roots

**Fig. S8.**
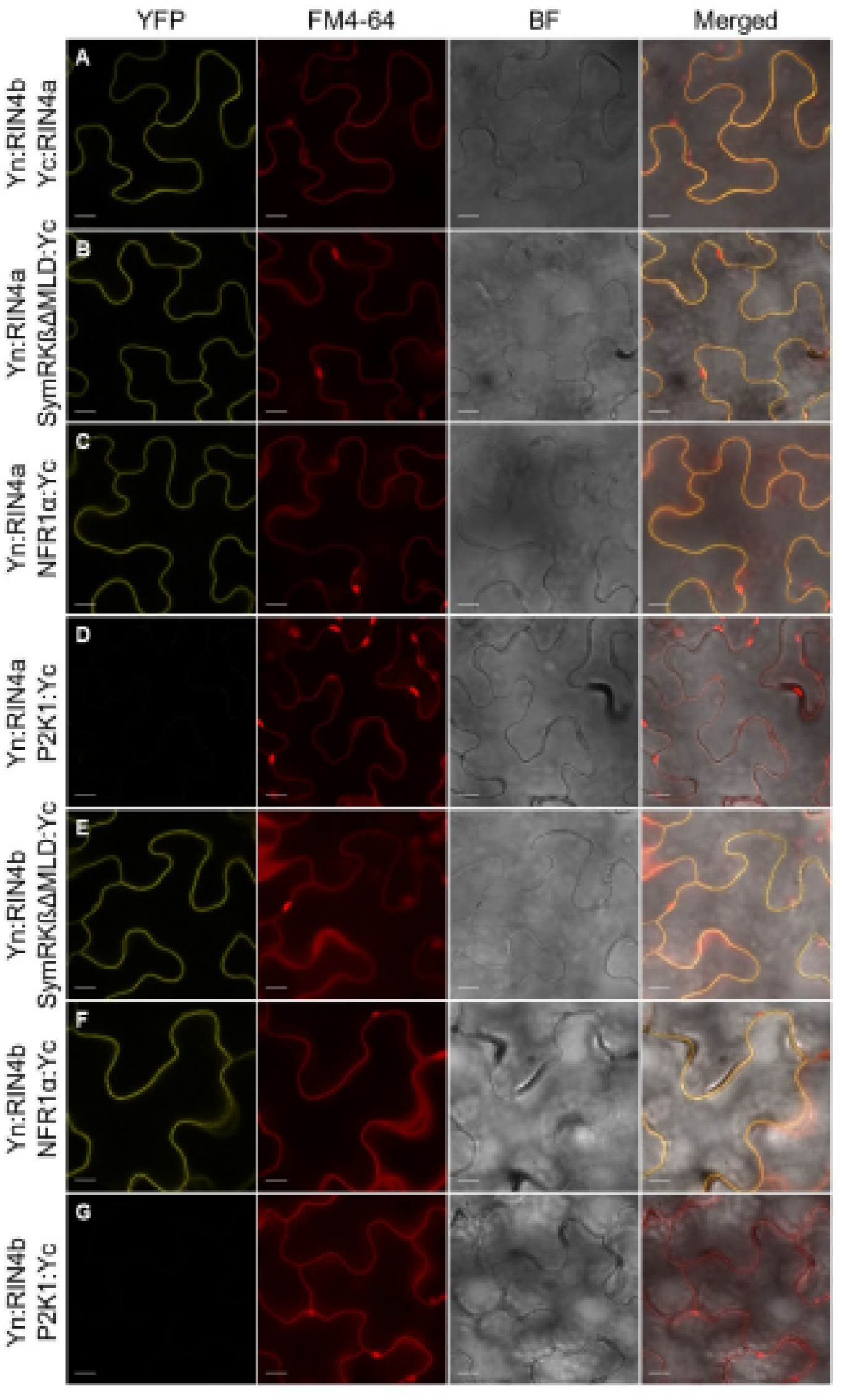
RIN4a and RIN4b interacts with symbiotic receptor-like kinases NFR1α and SymRKß in planta. (A) Bimolecular Fluorescence Complementation assay where the RIN4a and RIN4b interaction was used as a positive control. (B) RIN4a interacts with SymRKßΔMLD (C) and with NFR1α. (D) No fluorescence signal was observed when RIN4a was co-expressed with the Arabidopsis P2K1 receptor-like kinase, used as a negative control. (E) RIN4b interaction with SymRKßΔMLD (F) and with NFR1α. (G) No fluorescence signal was observed when RIN4b was co-expressed with P2K1. Left panels: BiFC, middle left panels FM4-64 staining of the plasma membrane (PM) to show that interaction occurs at the (PM). Middle right panels: bright field (BF). Right panels: merge of YFP (interaction signal) and red PM signal. Scale bars represent 10 μm. Leica SP8 confocal microscope was used to take images. At least three biological replicates were performed with similar results.

**Fig. S9.**
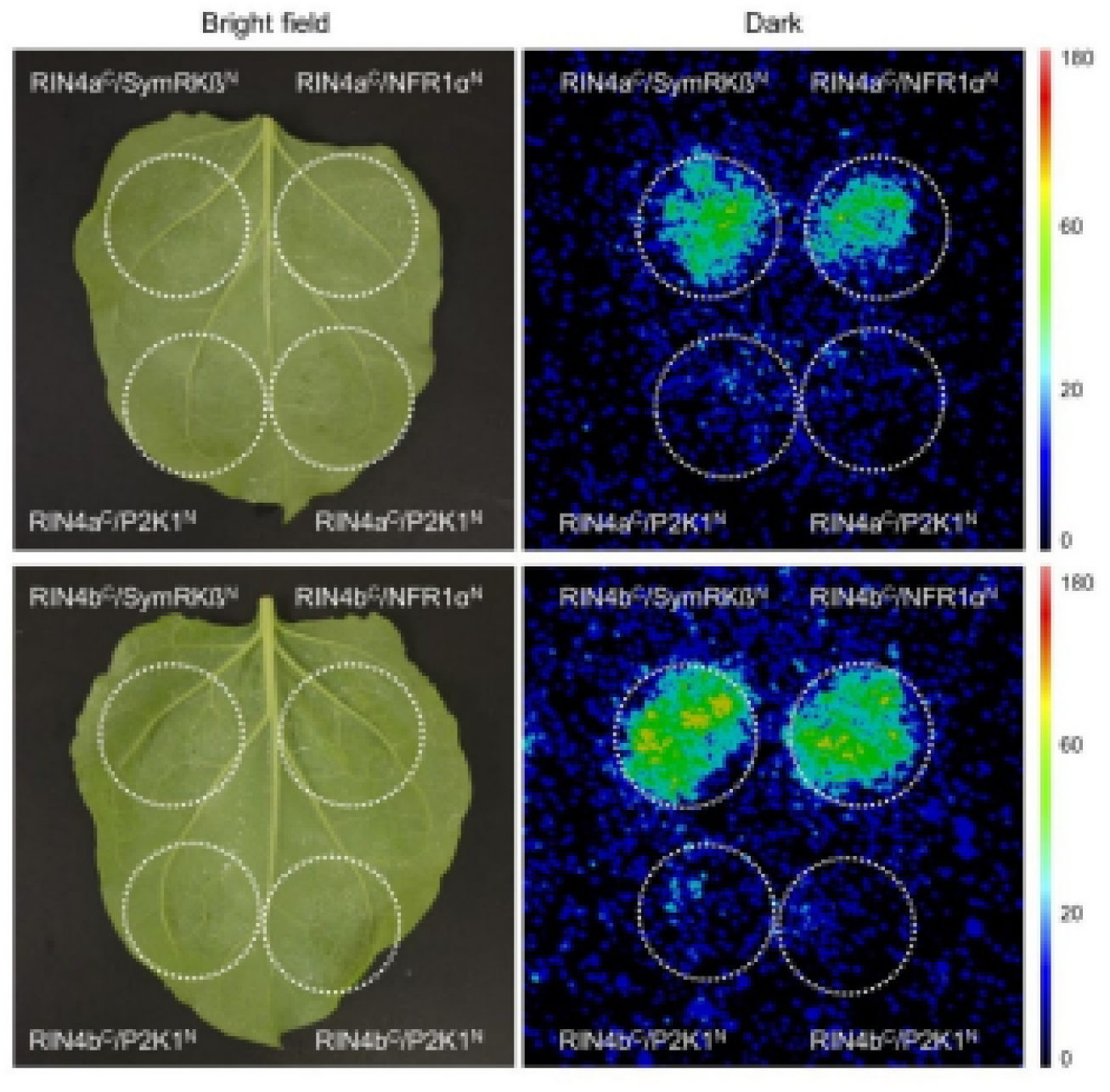
Split-Luciferase assay confirmed interaction of RIN4a and RIN4b with symbiotic receptors Upper panel shows RIN4a interaction with SymRKßΔMLD and NFR1α. Upon interaction between the co-expressed proteins fused to split domains of Luciferase, Luciferase activity is restored, and bioluminescence is detected (right panels). Lower panel shows RIN4b interaction with SymRKßΔMLD and NFR1α. In both cases Arabidopsis P2K1 receptor was used as negative control and no bioluminescence could be observed.

**Fig. S10.**
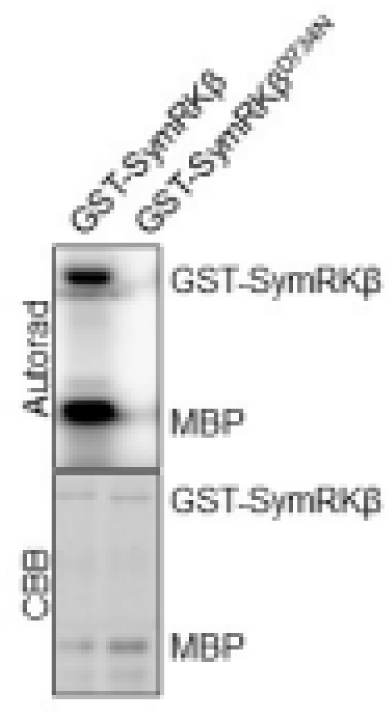
Soybean SymRKβ is an active kinase. Soybean SymRKβ kinase domain fused to GST was recombinantly expressed and kinase activity was detected *in vitro* using radioactive ATP. SymRKβ-KD trans-phosphorylates MBP substrate (left side). A point mutation was introduced to create a kinase inactive version, SymRKβ^D734N^, and it resulted in abolished kinase activity (right side, upper panel). Lower panel: CBB staining to show that both native and mutant versions were equally loaded.

**Fig. S11.**
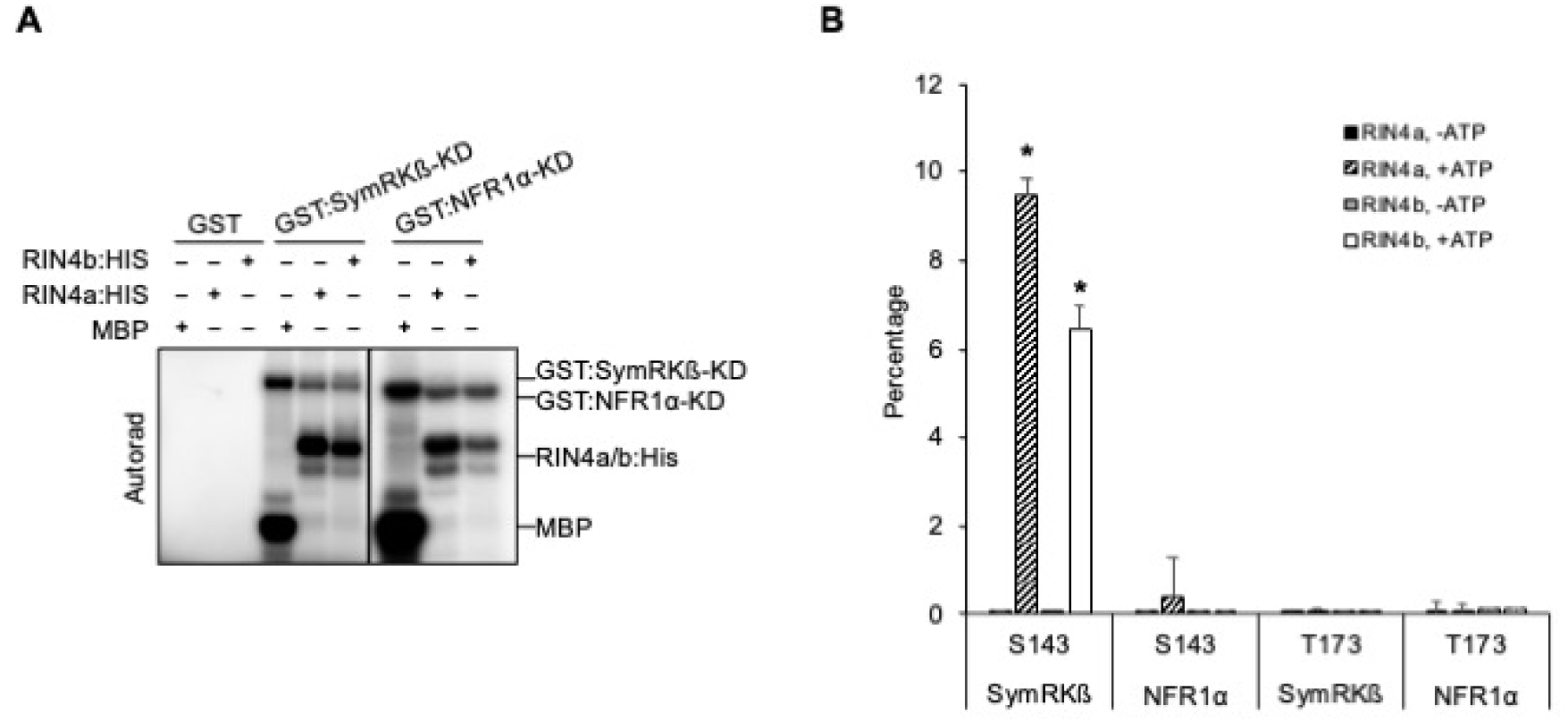
In vitro phosphorylation of nodulation-related S143 by SymRKβ. (A) Recombinantly expressed SymRKß and NFR1α kinase domains phosphorylate RIN4a and RIN4b *in vitro*. Myelin basic protein (MBP) was used as a positive control for SymRKβ and NFR1α kinase activity, and GST was used (left side) as a negative control. (B) Quantitative mass spectrometry (MS-MRM) was performed to identify the phosphorylation site triggered by the 2 kinases. S143 phosphorylation was phosphorylated only by SymRKß and quantified using heavy-labeled peptides generated against native peptides carrying S143. In both RIN4a and RIN4b S143 nodulation-related phosphorylation site is phosphorylated by SymRKβ whereas another phosphorylation site T173, used as negative control was not phosphorylated neither by SymRKß nor by NFR1α. MS run was performed in 2 biological and 3 technical replicates. Phosphorylation is expressed in percentage as the mean of all replicates. Error bars represent standard error. Student’s t-test * p<0.05

**Fig. S12.**
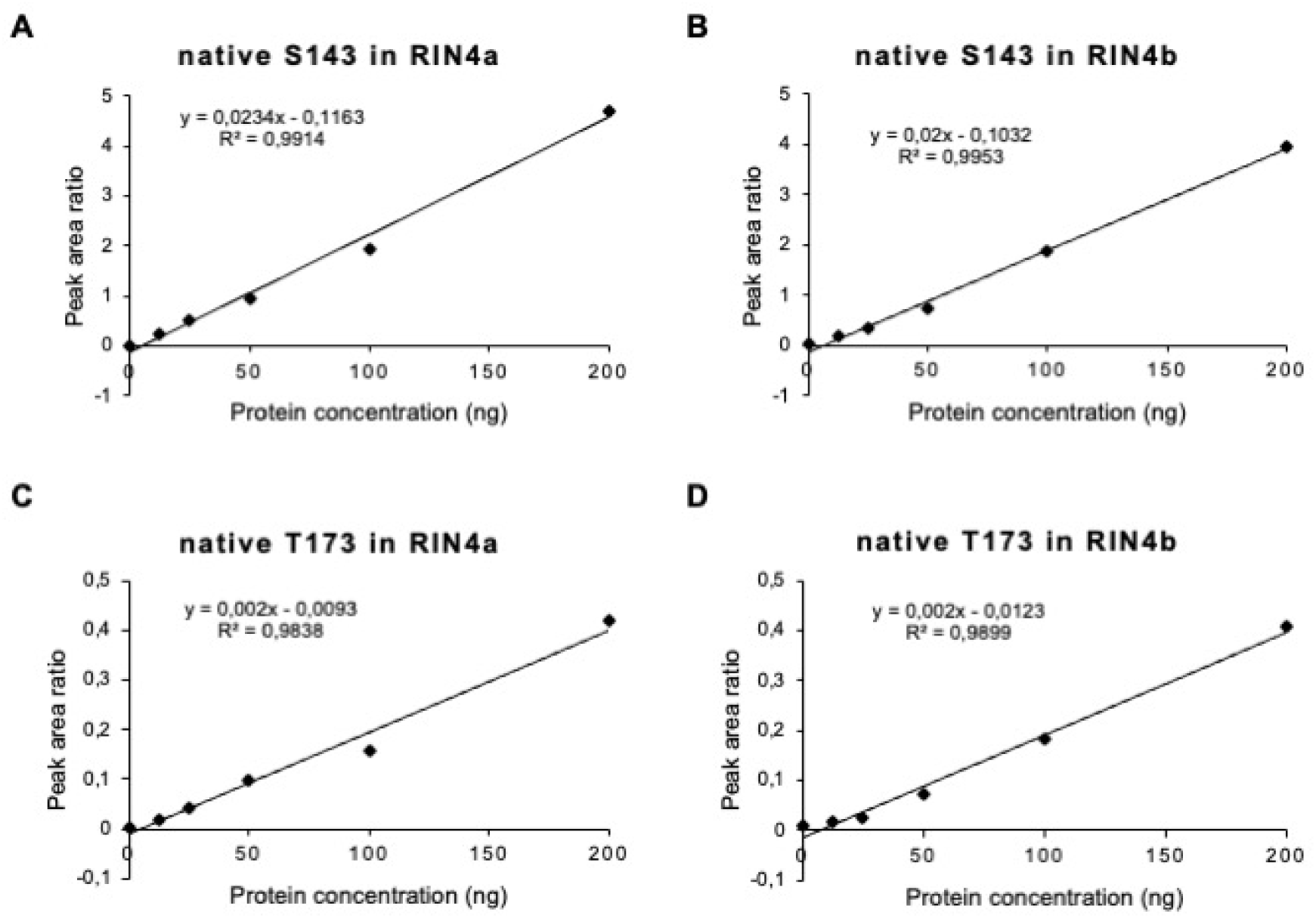
Linear relationship and correlation coefficients of RIN4a and RIN4b established for MS-SRM Calibration curves of S143 and T173 containing native peptides at different protein concentration of recombinantly expressed RIN4a and RIN4b proteins were established at fixed amount (100 fmol S143; 50 fmol T173) of heavy-labeled AQUA peptide. Both S143 and T173 carrying peptides are 100% identical between RIN4a and RIN4b.. The experiment was done in RIN4a and RIN4b recombinantly expressed protein background separately. Panels show a linear relationship between peak area ratio of native peptide normalized to heavy-labeled peptideand protein concentration . Scatter plots expressing the relationship between protein concentrations loaded per injection and peak area ratio normalized to heavy obtained per concentration gradient. Correlation coefficient (R^2^) values strongly support correlation between the protein concentration injected and the native peptide amount detected (normalized to heavy-labeled peptide).

**Table S1.** List of RIN4 proteins used for the phylogenetic tree.

| Full name<br>(Species) | Order |  | Abbreviation<br>Accession/Transcript ID | Note |
| --- | --- | --- | --- | --- |
| *Abrus precatorius* | Fabales | Apre027334769 | XP_027334769 |  |
| *Arachis ipaensis* | Fabales | AipaQ2FNA | Araip.Q2FNA |  |
|  |  | AipaT1NKW | Araip.T1NKW |  |
| *A. duranensis* | Fabales | Adur7K02I | Aradu.7K02I |  |
|  |  | Adur5N0SW | Aradu.5N0SW |  |
| *Cajanus cajan* | Fabales | Ccaj45821 | Cajcaj_Ccajan_45821 |  |
|  |  | Ccaj38676 | Cajcaj_Ccajan_38676 |  |
|  |  | Ccaj16754 | KYP62701 |  |
| *Chamaecrista fasciculata* | Fabales | Cfas4019S22266 | Chafas_Chafa4019S22266 |  |
|  |  | Cfas10196S13514 | Chafas_Chafa10196S13514 |  |
| *Cicer arietinum* | Fabales | Cari11744 | Ca_11744 |  |
|  |  | Cari22651 | Ca_22651 |  |
|  |  | Cari11699 | Ca_11699 |  |
| *Faidherbia albida* | Fabales | Falb00479g0034 | Faialb_Faial00479g0034.1 |  |
|  |  | Falb02280g0022 | Faialb_Faial02280g0022.1 |  |
| *Glycine max* | Fabales | Gma03G084000a | Glyma.03G084000.1 | RIN4a |
|  |  | Gma16G090700b | Glyma.16G090700.1 | RIN4b |
|  |  | Gma18G166800c | Glyma.18G166800.1 | RIN4c |
|  |  | Gma08G349500d | Glyma.08G349500.1 | RIN4d |
|  |  | Gma09G008700 | Glyma.09G008700.1 |  |
|  |  | Gma17G009600 | Glyma.17G009600.1 |  |
|  |  | Gma15G113100 | Glyma.15G113100.1 |  |
| *Lablab purpureus* | Fabales | Lpur000023g0028 | Labpur_Labpu000023g0028.1 |  |
|  |  | Lpur000066g0055 | Labpur_Labpu000066g0055.1 |  |
| *Lotus japonicus* | Fabales | Lj3g3v0730080 | Lj3g3v0730080.1 |  |
| *Lupinus angustifolius* | Fabales | Lang025313 | OIW18870 |  |
|  |  | Lang010450 | Lupang_Lup010450.1 |  |
|  |  | Lang012792 | Lupang_Lup012792.1 |  |
| *Medicago truncatula* | Fabales | Mtr8g012960 | Medtr8g012960.2 |  |
|  |  | Mtr7g056147 | Medtr7g056147.1 |  |
|  |  | Mtr2g025170 | Medtr2g025170.1 |  |
| *Mimosa pudica* | Fabales | Mpud10763S13484 | Mimpud_Mimpu10763S13484 |  |
| *Mucuna pruriens* | Fabales | Mpru76948 | Mucpru_lcl_QJKJ01009328.1_RDX76948.1_37901 |  |
|  |  | Mpru46803 | Mucpru_lcl_QJKJ01012707.1_RDX68051.1_46803 |  |
|  |  | Mpru58382 | Mucpru_lcl_QJKJ01017556.1_RDX58382.1_55230 |  |
| *Phaseolus vulgaris* | Fabales | Pvul010G021200 | Phvul.010G021200.1 |  |
|  |  | Pvul008G130600 | Phvul.008G130600.1 |  |
|  |  | Pvul006G144600 | Phvul.006G144600.1 |  |
| *Spatholobus suberectus* | Fabales | Ssub68649 | TKY68649 |  |
| *Trifolium pratense* | Fabales | Tpra17708 | Tp57577_TGAC_v2_mRNA17708 |  |
|  |  | Tpra6665 | Tp57577_TGAC_v2_mRNA6665 |  |
|  |  | Tpra13613 | Tp57577_TGAC_v2_mRNA13613 |  |
|  |  | Tpra25216 | Tp57577_TGAC_v2_mRNA25216 |  |
|  |  | Tpra24971 | Tp57577_TGAC_v2_mRNA24971 |  |
| *T. subterraneum* | Fabales | Tsub37391 | Trisub_lcl_DF974408.1_GAU48333.1_37391 |  |
|  |  | TsubDF973323 | Trisub_lcl_DF973323.1_GAU25<br>883.1_14941 |  |
| *Vigna*<br>*angularis* | Fabales | Vang08g04180 | XP_017442542 |  |
|  |  | Vang04g10640 | Vigang_vigan.Vang04g10640.1 |  |
|  |  | Vang03g07180 | Vigang_vigan.Vang03g07180.1 |  |
| *Vigna*<br>*radiata* | Fabales | Vrad09g08180 | A0A1S3VF94 |  |
|  |  | Vrad04g04050 | Vigrad_Vradi04g04050.1 |  |
|  |  | Vrad10g07710 | Vigrad_Vradi10g07710.1 |  |
| *V.*<br>*subterranean* | Fabales | Vsub107794g0016 | Vigsub_Vigsu107794g0016.1 |  |
|  |  | Vsub107732g0006 | Vigsub_Vigsu107732g0006.1 |  |
|  |  | Vsub107920g0110 | Vigsub_Vigsu107920g0110.1 |  |
| *V.*<br>*unguiculata* | Fabales | Vung10g046200 | Vigun10g046200.1 |  |
|  |  | Vung05g134100 | Vigun05g134100.1 |  |
|  |  | Vung06g156400 | Vigun06g156400.1 |  |
| *Chastanosper*<br>*mum*<br>*australe* | Fabales | Caus1227PA | Causus_Castanospermum21227-P<br>A |  |
|  |  | Caus26878PA | Causus_Castanospermum26878-P<br>A |  |
| *Cercis*<br>*canadensis* | Fabales | Ccan480S25307 | Cercan_Cerca480S25307 |  |
|  |  | Ccan86S33867 | Cercan_Cerca86S33867 |  |
| *Nissolia*<br>*schottii* | Fabales | Nsch1371S14229 | Nissch_Nissc1371S14229 |  |
|  |  | Nsch15112S15091 | Nissch_Nissc15112S15091 |  |
| *Parasponia*<br>*andersonii* | Rosales | PanPON36036 | PON36036 |  |
|  |  | PanPON52174 | PON52174 |  |
| *Trema orientale* | Rosales | TorPON59389 | PON59389 |  |
|  |  | TorPON91322 | PON91322 |  |
| *Dryas drummondii* | Rosales | Ddru380S21270 | Drydru_Drydr380S21270 |  |
| *Cannabis sativa* | Rosales | Csat030499662 | XP_030499662 |  |
| *Ceanothus thyrsiflorus* | Rosales | Cthy36985 | c36985_g1_i2 |  |
| *Fragaria vesca* | Rosales | Fves2g18650 | Fraves_FvH4_2g18650.1 |  |
| *Fragaria x ananassa* | Rosales | Fana00065658 | Fraana_FAN_iscf00065658.1.g00001.1 |  |
| *Humulus lupulus* | Rosales | HlupG023675 | Humlup_HL.SW.v1.0.G023675.1 |  |
|  |  | HlupG031954 | Humlup_HL.SW.v1.0.G031954.1 |  |
| *Morus notabilis* | Rosales | Mnot024026395 | XP_024026395 |  |
|  |  | Mnot013610 | EXB41536 |  |
| *Prunus persica* | Rosales | Pper8G199800 | Pruper_Prupe.8G199800.1 |  |
| *P. mume* | Rosales | Pmum008235453 | Prumum_lcl_NC_024131.1_XP_008235453.1_19524 |  |
|  |  | Pmum008224738 | Prumum_lcl_NC_024128.1_XP_008224738.1_9378 |  |
| *P. avium* | Rosales | Pavi0000558 | Pruavi_Pav_sc0000558.1_g920.1.mk |  |
| *Pyrus bretschneideri* | Rosales | Pbre009370675 | Pyrbre_lcl_NW_008988223.1_XP_009370675.1_22569 |  |
|  |  | Pbre009334905 | Pyrbre_lcl_NW_008988430.1_XP_009334905.1_32138 |  |
|  |  | Pbre009371881 | Pyrbre_lcl_NW_008988045.1_XP_009371881.1_2472 |  |
| *Rubus occidentalis* | Rosales | RoccG22517 | Rubocc_Bras_G22517 |  |
|  |  | RoccG10234 | Rubocc_Bras_G10234 |  |
| *Rosa chinensis* | Rosales | RchiChr6g0284211 | A0A2P6PU72 |  |
| *Ziziphus jujube* | Rosales | Zjuj015892585 | Zizjuj_lcl_NC_029687.1_XP_015892585.1_24446 |  |
|  |  | Zjuj015887188 | Zizjuj_lcl_NC_029685.1_XP_015887188.1_19239 |  |
|  |  | Zjuj024926926 | Zizjuj_lcl_NC_029679.1_XP_024926926.1_1025 |  |
| *Alnus glutinosa* | Fagales | Agl60713S33275 | Alngl60713S33275 |  |
|  |  | Agl92815S37678 | Alngl92815S37678 |  |
| *Casuarina glauca* | Fagales | Cgl230S11910 | Casgl230S11910 |  |
|  |  | Cgl46S18253 | Casgl46S18253 |  |
| *Junglans regia* | Fagales | Jreg38502 | Jugreg_g38502.t1 |  |
|  |  | Jreg49802 | Jugreg_g49802.t1 |  |
|  |  | Jreg27489 | Jugreg_g27489.t1 |  |
|  |  | Jreg5419 | Jugreg_g5419.t1 |  |
|  |  | Jreg8286 | Jugreg_g8286.t1 |  |
| *Quercus robur* | Fagales | QrobP0336020 | Querob_Qrob_P0336020.2 |  |
| *Q. lobata* | Fagales | Qlob030970650 | XP_030970650 |  |
| *Datisca glomerata* | Cucurbitales | Dgl27393S26490 | Datgl27393S26490 |  |
|  |  | Dgl216S24577 | Datgl216S24577 |  |
|  |  | Dgl1602S21798 | Datgl1602S21798 |  |
| *Begonia fuchsioides* | Cucurbitales | Bfuc8283S41985 | Begfuc_Begfu8283S41985 |  |
|  |  | Bfuc141S19375 | Begfuc_Begfu141S19375 |  |
|  |  | Bfuc5443S37656 | Begfuc_Begfu5443S37656 |  |
|  |  | Bfuc91277S43551 | Begfuc_Begfu91277S43551 |  |
|  |  | Bfuc719S40210 | Begfuc_Begfu719S40210 |  |
|  |  | Bfuc646S39458 | Begfuc_Begfu646S39458 |  |
|  |  | Bfuc2878S29171 | Begfuc_Begfu2878S29171 |  |
| *Citrullus lanatus* | Cucurbitales | Clan014936 | Citlan_Cla014936 |  |
|  |  | Clan021409 | Citlan_Cla021409 |  |
|  |  | Clan004570 | Citlan_Cla004570 |  |
| *Cucumis sativus* | Cucurbitales | Cusa078160 | Cucsa.078160.1 |  |
|  |  | Cusa256560 | Cucsa.256560.2 |  |
|  |  | CusaChr13333 | XP_011660296 |  |
| *Cucumis melo* | Cucurbitales | Cmel3C021057 | XP_008458026 |  |
|  |  | CmelC006302 | XP_008438343 |  |
|  |  | Cmel3C021706 | Cucmel_MELO3C021706.2.1 |  |
| *Cucurbita pepo* | Cucurbitales | CpepLG12g11050 | Cucpep_Cp4.1LG12g11050.1 |  |
|  |  | Cpep17g11110 | Cucpep_Cp4.1LG17g11110.1 |  |
|  |  | Cpep1LG03g18010 | Cucpep_Cp4.1LG03g18010.1 |  |
| *Cucurbita maxima* |  | CmaxCh14G019030 | Cucmax_CmaCh14G019030.1 |  |
|  |  | CmaxCh06G012380 | Cucmax_CmaCh06G012380.1 |  |
| *Cucurbita moschata* |  | CmosCh08G002240 | XP_022953250 |  |
|  |  | CmosCh17G012410 | Cucmos_CmoCh17G012410.1 |  |
|  |  | CmosCh14G019560 | Cucmos_CmoCh14G019560.1 |  |
| *Lagenaria siceraria* | Cucurbitales | Lsic05G018530 | Lagsic_Lsi05G018530.1 |  |
|  |  | Lsic02G010850 | Lagsic_Lsi02G010850.1 |  |
| *Momordica charantia* | Cucurbitales | Mcha022145992 | XP_022145992 |  |
|  |  | Mcha022142617 | Momcha_lcl_NW_019104509.1<br>_XP_022142617.1_13621 |  |
|  | Non<br>FaFaCu<br>Ro |  |  |  |
| *Arabidopsis thaliana* | Non<br>FaFaCu<br>Ro | RIN4<br>AT3G25070 | AT3G25070 | AtRIN4 |
| *A. lyrata* | Non<br>FaFaCu<br>Ro | Alyr3G40780 | AL3G40780.t1 |  |
| *Brassica rapa* | Non<br>FaFaCu<br>Ro | BraC03983 | Brara.C03983.1 |  |
| *Brachypodium distachyon* | Non<br>FaFaCu<br>Ro | Bdi1g01730 | Bradi1g01730.3 |  |
|  |  | Bdi3g40950 | Bradi3g40950.1 |  |
| *Setaria viridis* | Non<br>FaFaCu<br>Ro | Svir6G226900 | Sevir.6G226900.1 |  |
|  |  | Svir9G012200 | Sevir.9G012200.1 |  |
| *Zea mays* | Non<br>FaFaCu<br>Ro | GRMZM2G012229 | GRMZM2G012229_T01 |  |
|  |  | GRMZM2G703858 | GRMZM2G703858_T02 |  |
| *Oryza sativa* | Non<br>FaFaCu<br>Ro | Os03g63140 | LOC_Os03g63140.1 |  |
|  |  | Os08g41470 | LOC_Os08g41470.1 |  |
| *Nicotiana benthamiana* | Non FaFaCu Ro | NbenAPY20266 | APY20266 |  |
| *N. silvestris* | Non FaFaCu Ro | NsilA0A1S3ZGX8 | A0A1U7YR47 |  |
| *Solanum lycopersicum* | Non FaFaCu Ro | Slyc09g059430 | Solyc09g059430.2.1 |  |
|  |  | Slyc06g083390 | Solyc06g083390.2.1 |  |
|  |  | Slyc11g012010 | Solyc11g012010.2.1 |  |

**Table S2:** Summary of *L. japonicus rin4* mutant screen.

| Mutant ID | Total No. of plants | Wild type | Heterozygous | Homozygous |
| --- | --- | --- | --- | --- |
| 30019656 | 8 | 3 | 5 | 0 |
| 30000711 | 10 | 3 | 7 | 0 |

**Table S3.** Soybean RIN4 native and AQUA peptides.

| Peptide name | Native peptide sequence | Heavy-labeled peptide sequence | Note |
| --- | --- | --- | --- |
| S143 (non phosphorylated) | APGRDSPQWEPK | R.APGRDSPQWEP.[K_C13N15] | RIN4a and RIN4b sequences are identical |
| pS143 (phosphorylated) | APGRD(pS)PQWEPK | R.APGRD(pS)PQWEP.[K_C13N15] | RIN4a and RIN4b sequences are identical |
| T173 (non phosphorylated) | GDETPDKGAAVPK | R.GDETPDKGAAVP.[K_C13N15] | RIN4a and RIN4b sequences are identical |
| pT173 (phosphorylated) | GDE(pT)PDKGAAVPK | R.GDE(pT)PDKGAAVP.[K_C13N15] | RIN4a and RIN4b sequences are identical |

## References

1. Oldroyd GE, et al. (2011) The rules of engagement in the legume-rhizobial symbiosis. Annu Rev Genet. 45:119–44.

2. Quilbé J, et al. (2022) Molecular Mechanisms of Intercellular Rhizobial Infection: Novel Findings of an Ancient Process. Front. Plant Sci. 13. doi: 10.3389/fpls.2022.922982

3. Broghammer A, et al. (2012) Legume receptors perceive the rhizobial lipochitin oligosaccharide signal molecules by direct binding. Proc Natl Acad Sci U S A. 109(34):13859–64.

4. Radutoiu S, et al. (2003) Plant recognition of symbiotic bacteria requires two LysM receptor-like kinases. Nature. 425(6958):585–92.

5. Indrasumunar A, et al. (2010) Inactivation of duplicated nod factor receptor 5 (NFR5) genes in recessive loss-of-function non-nodulation mutants of allotetraploid soybean (Glycine max L. Merr.). Plant Cell Physiol. 51(2):201–14.

6. Indrasumunar A, et al. (2011). Nodulation factor receptor kinase 1α controls nodule organ number in soybean (Glycine max L. Merr). Plant J. 65(1):39–50.

7. Stracke S, et al. (2002) A plant receptor-like kinase required for both bacterial and fungal symbiosis. Nature. 417(6892):959–62.

8. Indrasumunar A, et al. (2015) Functional analysis of duplicated Symbiosis Receptor Kinase (SymRK) genes during nodulation and mycorrhizal infection in soybean (Glycine max). J Plant Physiol. 176:157–68.

9. Gherbi H, et al. (2008) SymRK defines a common genetic basis for plant root endosymbioses with arbuscular mycorrhiza fungi, rhizobia, and Frankiabacteria. Proc Natl Acad Sci U S A. 105(12):4928–32.

10. Antolín-Llovera M, Ried MK, Parniske M (2014) Cleavage of the SYMBIOSIS RECEPTOR-LIKE KINASE ectodomain promotes complex formation with Nod factor receptor 5. Curr Biol. 24:422–7.

11. Madsen EB, et al. (2011) Autophosphorylation is essential for the in vivo function of the Lotus japonicus Nod factor receptor 1 and receptor-mediated signalling in cooperation with Nod factor receptor 5. Plant J. 65(3):404–17.

12. Murakami E, et al. (2018) Epidermal LysM receptor ensures robust symbiotic signalling in Lotus japonicus. Elife. pii: e33506.

13. Wong JEMM, et al. (2019) A Lotus japonicus cytoplasmic kinase connects Nod factor perception by the NFR5 LysM receptor to nodulation. Proc Natl Acad Sci U S A. 14339–14348.

14. Singh S, et al. (2014) CYCLOPS, a DNA-binding transcriptional activator, orchestrates symbiotic root nodule development. Cell Host Microbe. 15(2):139–52.

15. Soyano T, et al. (2013) Nodule inception directly targets NF-Y subunit genes to regulate essential processes of root nodule development in Lotus japonicus. PLoS Genet. 9(3):e1003352.

16. Serna-Sanz A, Parniske M, Peck SC (2011) Phosphoproteome analysis of Lotus japonicus roots reveals shared and distinct components of symbiosis and defense. Mol Plant Microbe Interact. 24(8):932–7.

17. Nguyen TH, et al. (2012) Quantitative phosphoproteomic analysis of soybean root hairs inoculated with Bradyrhizobium japonicum. Mol Cell Proteomics. 11(11):1140–55.

18. Rose CM, et al. (2012) Rapid Phosphoproteomic and Transcriptomic Changes in the Rhizobia-legume Symbiosis. Mol Cell Proteomics. 11(9): 724–744.

19. Roy S, et al. (2019) Celebrating 20 years of genetic discoveries in legume nodulation and symbiotic nitrogen fixation. Plant Cell. pii: tpc.00279.2019.

20. Gage DJ, (2004) Infection and invasion of roots by symbiotic, nitrogen-fixing rhizobia during nodulation of temperate legumes. Microbiol Mol Biol Rev. 68(2):280–300.

21. Mackey D, et al. (2002) RIN4 interacts with Pseudomonas syringae type III effector molecules and is required for RPM1-mediated resistance in Arabidopsis. Cell. 108(6):743–54.

22. Ray SK, et al. (2019) Role of RIN4 in Regulating PAMP-Triggered Immunity and Effector-Triggered Immunity: Current Status and Future Perspectives. Mol Cells 42(7):503–511.

23. Mackey D, et al. (2003) Arabidopsis RIN4 is a target of the type III virulence effector AvrRpt2 and modulates RPS2-mediated resistance. Cell. 112(3):379–89.

24. Toruño TY, Shen M, Coaker G, Mackey D (2019) Regulated Disorder: Posttranslational Modifications Control the RIN4 Plant Immune Signaling Hub. Mol Plant Microbe Interact. 32(1):56–64.

25. Sun X, et al. (2014) The intrinsically disordered structural platform of the plant defence hub protein RPM1-interacting protein 4 provides insights into its mode of action in the host-pathogen interface and evolution of the nitrate-induced domain protein family. FEBS J. 281(17):3955–79.

26. Lee D, et al. (2015) Phosphorylation of the Plant Immune Regulator RPM1- INTERACTING PROTEIN4 Enhances Plant Plasma Membrane H⁺-ATPase Activity and Inhibits Flagellin-Triggered Immune Responses in Arabidopsis. Plant Cell. 27:2042–56.

27. Darling AL, Uversky VN (2018) Intrinsic Disorder and Posttranslational Modifications: The Darker Side of the Biological Dark Matter. Front Genet. 9:158.

28. Kim MG, et al. (2005) Two Pseudomonas syringae type III effectors inhibit RIN4- regulated basal defense in Arabidopsis. Cell. 121:749–59.

29. Chung EH, et al. (2014) A Plant Phosphoswitch Platform Repeatedly Targeted by Type III Effector Proteins Regulates the Output of Both Tiers of Plant Immune Receptors. Cell Host & Microbe 16: 484–494.

30. Redditt TJ, et al. (2019) AvrRpm1 Functions as an ADP-Ribosyl Transferase to Modify NOI Domain-Containing Proteins, Including Arabidopsis and Soybean RPM1-Interacting Protein4. Plant Cell. 31: 2664–2681.

31. Liu J, et al. (2011) A receptor-like cytoplasmic kinase phosphorylates the host target RIN4, leading to the activation of a plant innate immune receptor. Cell Host Microbe. 9(2):137–46.

32. Xu N, et al. (2017) The Bacterial Effector AvrB-Induced RIN4 Hyperphosphorylation Is Mediated by a Receptor-Like Cytoplasmic Kinase Complex in Arabidopsis. Mol Plant Microbe Interact. 30(6):502–512.

33. Selote D and Kachroo A (2010) RPG1-B-Derived Resistance to AvrB-expressing *Pseudomonas syringae* requires RIN4-like proteins in Soybean. Plant Physiology 153(3): 1199–1211.

34. Parniske M (2018) Uptake of bacteria into living plant cells, the unifying and distinct feature of the nitrogen-fixing root nodule symbiosis. Curr Opin Plant Biol. 44:164–174.

35. Griesmann M, et al. (2018). Phylogenomics reveals multiple losses of nitrogen-fixing root nodule symbiosis. Science. 361(6398).

36. Afzal AJ, Kim JH, Mackey D (2013) The role of NOI-domain containing proteins in plant immune signaling. BMC Genomics. 14:327.

37. Li W, Lan P (2015) Re-analysis of RNA-seq transcriptome data reveals new aspects of gene activity in Arabidopsis root hairs. Front Plant Sci 6:421.

38. Małolepszy A, et al. (2016) The LORE1 insertion mutant resource. Plant J 88(2):306–317.

39. Mun T, et al. (2016) Lotus Base: An integrated information portal for the model legume Lotus japonicus. Sci Rep 6:39447.

40. Michno JM, et al. (2020) Integration, abundance, and transmission of mutations and transgenes in a series of CRISPR/Cas9 soybean lines. BMC Biotechnol 20:10.

41. Choudhury SR, Pandey S (2015) Phosphorylation-Dependent Regulation of G-Protein Cycle during Nodule Formation in Soybean. Plant Cell 27: 3260–3276.

42. Yoshida S and Parniske M (2005) Regulation of plant symbiosis receptor kinase through serine and threonine phosphorylation. J Biol Chem 280(10):9203–9.

43. Kettenbach AN, Rush J, Gerber SA (2011) Absolute quantification of protein and post translational modification abundance with stable isotope-labeled synthetic peptides. Nat Protoc. 175–86.

44. Schauser L, et al. (1999) A plant regulator controlling development of symbiotic root nodules. Nature. 402(6758):191–5.

45. Cerri MR, et al. (2017) The ERN1 transcription factor gene is a target of the CCaMK/CYCLOPS complex and controls rhizobial infection in Lotus japonicus. New Phytol. 215(1):323–337.

46. Hayashi S, et al. (2012) Transient Nod factor-dependent gene expression in the nodulation-competent zone of soybean (Glycine max [L.] Merr.) roots. Plant Biotechnol J. 10(8):995–1010.

47. Kawaharada Y, et al. (2017) The Ethylene Responsive Factor Required for Nodulation 1 (ERN1) Transcription Factor Is Required for Infection-Thread Formation in *Lotus japonicus*. Mol Plant Microbe Interact. 30(3):194–204.

48. Singh S, et al. (2014) CYCLOPS, A DNA-Binding Transcriptional Activator, Orchestrates Symbiotic Root Nodule Development. Cell Host & Microbe 15, 139–152.

49. Cao Y, Halane MK, Gassmann W, Stacey G. (2017) The Role of Plant Innate Immunity in the Legume-Rhizobium Symbiosis. Annu Rev Plant Biol. 68:535–561.

50. Tóth K, Stacey G (2015) Does plant immunity play a critical role during initiation of the legume-rhizobium symbiosis? Front Plant Sci. 6:401.

51. Chuberre C, et al. (2018) Plant Immunity Is Compartmentalized and Specialized in Roots. Front Plant Sci 9:1692.

52. Okazaki S, et al. (2013) Hijacking of leguminous nodulation signaling by the rhizobial type III secretion system. Proc Natl Acad Sci U S A. 10(42):17131–6.

53. Wagner MR, et al. (2016) Host genotype and age shape the leaf and root microbiomes of a wild perennial plant. Nat. Commun. 7:12151 doi: 10.1038/ncomms12151.

## References

1. Salgado MG, et al. (2018) Comparative Analysis of the Nodule Transcriptomes of Ceanothus thyrsiflorus (Rhamnaceae, Rosales) and Datisca glomerata (Datiscaceae, Cucurbitales). Front Plant Sci. 9:1629.

2. Leebens-Mack JH, et al (2019) One thousand plant transcriptomes and the phylogenomics of green plants. Nature 574(7780):679–685.

3. Saitou N, Nei M (1987) The neighbor-joining method: a new method for reconstructing phylogenetic trees. Mol Biol Evol. 1987;4: 406–425.

4. Jukes TH (2000) The Neutral Theory of Molecular Evolution. Genetics 2000;154: 956–958.

5. Tóth K, Batek J, and Stacey G (2016) Generation of Soybean (Glycine max) Transient Transgenic Roots. Curr. Protoc. Plant Biol. 1–13.

6. Michno JM, et al. (2015) CRISPR/Cas mutagenesis of soybean and Medicago truncatula using a new web-tool and a modified Cas9 enzyme. GM Crops Food. 6(4):243–52.

7. Curtin SJ, et al. (2011) Targeted mutagenesis of duplicated genes in soybean with zinc finger nucleases. Plant Physiol. 156(2):466–73.

8. Konieczny A, Ausubel FM (1993) A procedure for mapping Arabidopsis mutations using co-dominant ecotype-specific PCR-based markers. Plant J. 4(2):403–10.

9. Jacobs TB, LaFayette PR, Schmitz RJ, Parrott WA (2015) Targeted genome modifications in soybean with CRISPR/Cas9. BMC Biotechnol. 15:16.

10. Zhu X, et al. (2014). An efficient genotyping method for genome-modified animals and human cells generated with CRISPR/Cas9 system. Sci Rep. 4:6420.

11. Li H, et al. (2010) Misexpression of miR482, miR1512, and miR1515 increases soybean nodulation. Plant Physiol. 153(4):1759–70.

12. Graham TL, Graham MY, Subramanian S, Yu O (2007) RNAi silencing of genes for elicitation or biosynthesis of 5-deoxyisoflavonoids suppresses race-specific resistance and hypersensitive cell death in Phytophthora sojae infected tissues. Plant Physiol. 144(2):728–40.

13. Lefebvre B, et al. (2010) A remorin protein interacts with symbiotic receptors and regulates bacterial infection. Proc Natl Acad Sci 107(5):2343–8.

14. Cho SH, et al. (2022) Activation of the plant mevalonate pathway by extracellular ATP. Nat Commun 13(1):450.

15. Libault M, et al. (2008) Identification of four soybean reference genes for gene expression normalization. Plant Genome 1: 44–54.

16. Schmittgen TD, Livak KJ (2008) Analyzing real-time PCR data by the comparative C(T) method. Nat Protoc. 3(6):1101–8.

17. Untergasser A, et al. (2012). Primer3--new capabilities and interfaces. Nucleic Acids Res. 40(15):e115.

